# monitoraSom: an easy path from Soundscapes to Ecology

**DOI:** 10.1101/2025.06.23.661148

**Authors:** G. L. M. Rosa, J. P. Zurano, I. M. D. Torres, C. R. Simoes, L. dos Anjos, C. B. de Araujo

## Abstract

Soundscape analysis for biodiversity monitoring with data from autonomous recorders has become a vital tool for biodiversity research. Yet, workflows fragmented across multiple software and technical barriers make the analysis of such large datasets a challenging task. Here, we present ’monitoraSom’, an efficient and user-friendly toolbox that facilitates the path between Passive Acoustic Monitoring (PAM) with ecological studies. This R package includes (1) modular workflows adaptable to diverse study designs, (2) an efficient and responsive user interface for fast soundscape recording segmentation; (3) validation methods adaptable to most data availability scenarios, including an interface for manual validation, and (4) dynamic diagnostics for threshold optimization to refine detection performance. The package is designed for scalability, with simple and easy to use batch processing and parallelization options, as well as simple data formats, making it accessible to beginners and advanced users. By reducing the technical hurdles and ensuring reproducibility, it accelerates the transformation of acoustic data into ecological information from rare species monitoring to community level surveys.

## 1. INTRODUCTION

Passive Acoustic Monitoring (PAM) has emerged as a pivotal tool in ecological research, driven by advancements in autonomous recording units (ARU) that enable large scale and cost-effective sampling of sound producing organisms (Hoefer et al. 2023). New venues for research are opening, and it is becoming a critical asset in the race to keep pace with the rapid habitat degradation and climate change (Desjonqueres et al. 2022). Also, the possibility of lossless storage in digital repositories further enhances the value of PAM, allowing researchers to revisit the data for new insights (Ross et al. 2023). Despite these opportunities, analyzing large acoustic datasets is still challenging due to the sheer volume of data and workflow complexity. Recording soundscapes over extended periods generates terabytes of data, requiring robust storage solutions and high-performance processing capabilities (Napier et al 2024). Although considerable strides were made in the development of computationally efficient tools, programming ability is still necessary to solve the compound inefficiencies and interoperability issues of a complete PAM workflow.

The segmentation of soundscape recordings to find and register temporal and spectral regions of interest (ROI) that contain the signals of the target organisms is usually one the most time consuming and critical phases of the PAM workflow. Although unsupervised segmentation (Ulloa et al. 2018) could reduce manual effort, manual segmentation is still essential for detection benchmarking. Thus, observers capable of accurately discerning and correctly labeling the acoustic signals of the studied taxa, a critical asset for PAM research, are burdened with cumbersome file management and disjoint workflows. A simplified and locally adaptable workflow is crucial for increasing PAM’s efficiency and promoting its broader adoption.

Here we introduce ’monitoraSom’, an R package designed to overcome these challenges and unify the PAM workflow within a user-friendly framework. The key contributions of ’monitoraSom’ are integrated tools for segmentation, validation, and performance evaluation operable entirely within R. Among its core functionalities, we highlight the graphical interfaces designed to accelerate manual segmentation of soundscape recordings and detection validation. Additionally, ’monitoraSom’ includes dedicated tools for validation diagnostics, offering quantitative assessments of detection performance. The modular design of the functions supports creative customization, encourages methodological experimentation, and simplifies the processing and analysis of large volumes of data generated by PAM. By consolidating these features, ’monitoraSom’ reduces the learning curve associated with scripting and empowers users to focus on ecological questions rather than technical hurdles, accelerating the transformation of raw acoustic data into actionable knowledge.

## 2. PACKAGE OVERVIEW

### a. PAM workflow overview

While ’monitoraSom’ is modular and adaptable to diverse analytical pipelines, we outline a simplified workflow to detect acoustic signals of interest (Figure **1**). Before starting the analysis, users must set a local working directory for the project and populate it with the adequate subdirectories to receive the data by using the ’set_workspace’ function (see section 3 Installation and setup).

**Figure 1:**
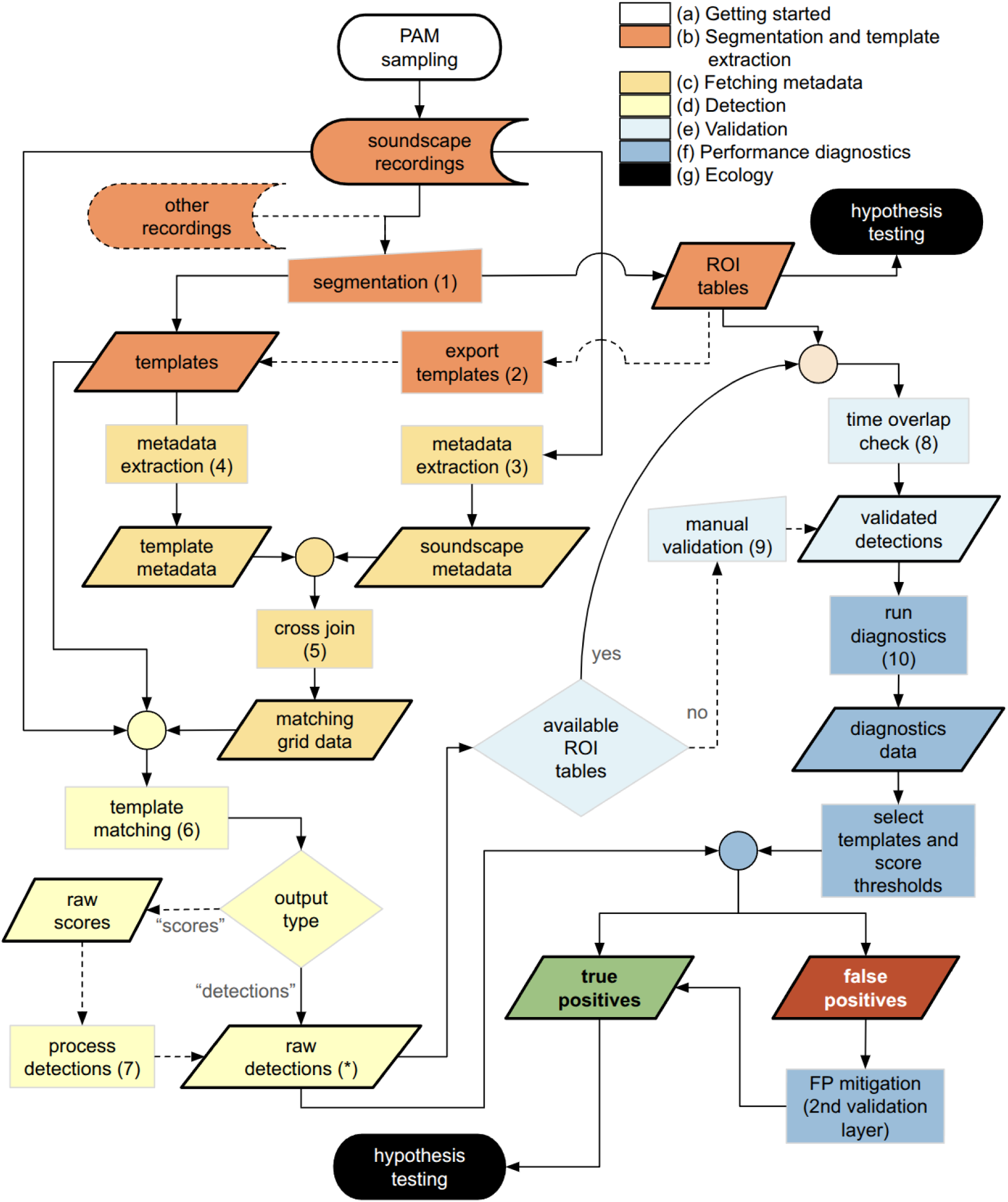
Flowchart of the ’monitoraSom’ workflow. The numbers indicate the functions dedicated to each respective step as follows: (1) ’launch_segmentation_app’; (2) ’export_roi_cuts’; (3) ’fetch_soundscape_metadata’; (4) ’fetch_template_metadata’; (5) ’fetch_match_grid’; (6) ’run_matching’; (7) ’fetch_score_peaks’; (8) ’validate_by_overlap’; (9) ’launch_validation_app’; (10) ’diagnostic_validations’.

The workflow begins with spectrogram segmentation, where users identify and label regions of interest (ROI) containing signals from target organisms (see 4 segmentation app). Segmentation involves characterizing the sounds within a recording by highlighting their temporal position and frequency band. To streamline this foundational step, we developed a dedicated interface launched via ’launch_segmentation_app’ function. The key features of the segmentation app include: automated file management to ensure consistent input and output; preset species lists to accelerate ROI labelling; and a streamlined and responsive interface for quick navigation within and between soundscapes. Within the app, users can generate templates by exporting selected ROls externally to R as WAV files, either from soundscapes or recordings from other sources, enabling interactive control over downstream steps of the detection workflow. Corresponding ROI tables are automatically generated for each segmented soundscape recording and stored as standardized CSV files for downstream use.

The next step is the automated signal detection using template matching. In template matching two sets of data are used, one or more searching templates, and the set of soundscapes. The process begins by importing the data that will guide the analyses (see 5 Fetching metadata). First, metadata consistency of soundscapes and templates is verified using ’fetch_soundscape_metadata’ and ’fetch_template_metadata’. Next, the template-soundscape compatibility is checked to generate a search grid with the function ’fetch_match_grid’, which can be further processed for customization of the analysis (e.g., removing pairs between templates and their soundscape recordings of origin or customizing spectral parameters). Next, the ’run_matching’ function uses the search grid to compute cross-correlations that will determine a detection. Detections obtained with high correlation scores will be considered the best candidates to be true occurrences of the target signals. Although this part of the ’monitoraSom’ workflow is designed to allow granular control and customization at multiple points of the analysis, we have implemented alternatives for simplified use cases (see section 6 Detection) With detections at hand, the next step is to validate them as true/false positives (TP/FP) and identify false negatives (FN). When reference ROI tables are available for enough soundscapes, a *priori* validation can be automatically computed by checking for temporal overlap between detections and ROls labeled as belonging to the same target species with the function ’validate_by_overlap’. ROI tables can be imported back to the R session on batch by using the ’fetch_rois’ function. If ROI tables are not available, the validation app (function ’launch_validation_app’) enables manual validation a *posteriori* via side-by-side comparison between templates and detections. It should be noted that the a *priori* validation with segmentation data is preferable over the a *posteriori* alternative because it includes the assessment of false negatives, which is critical for measuring detection performance. While flexible, the a *posteriori* approach does not account for false negatives and can be prone to observer biases, thus we recommend reserving it for exploratory analyses or validation under severely limited data availability (see Supplementary Material S1 for best practices).

Validation results feed directly into ’monitoraSom’ performance evaluation tools, in which each template is evaluated as a single-class classification model, with performance assessed across a user-defined range of score thresholds (see Balantic & Donovan 2020). Key diagnostic metrics enable data driven optimization of template performance through score threshold selection. In other words, users can fine tune the detection performance to specific goals, such as rare species monitoring or community level surveys. With this iterative refining process, users can distill raw detections into biologically meaningful data, ready for downstream analysis and biological hypothesis testing.

### b. Design principles

The objective of ’monitoraSom’ is to provide a user-friendly and versatile toolkit for bioacoustic monitoring and analysis. Intuitive interface and streamlined commands enable users to go from raw recordings to a set of validated detections without the need of advanced coding skills. Also, the automation of batch processing steps reduces the workload for experiment design customization. Function modularity and output formats have a clear organization to facilitate adaptability and extensibility with third-party software for integration with existing workflows. For instance, the segmentation app can be seamless integration into workflows beyond the traditional objectives of PAM, such as signal processing, acoustic measurement, and general sound analysis. Additionally, after minimal input format adaptation the validation app allows can be useful to validate detections generated by other software (e.g., BirdNet), improving the interoperability with other existing detection workflows.

## 3. The workflow of ’monitoraSom’

### a. Getting started

Currently the ’monitoraSom’ package can be installed from within an R session using the following commands:

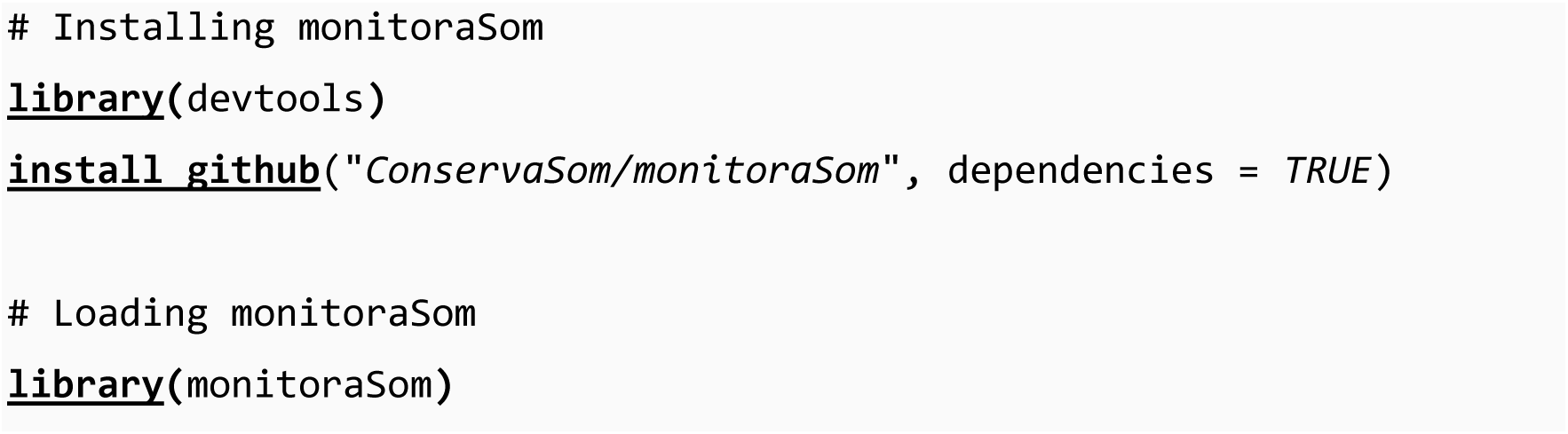

The ’monitoraSom’ workflow, particularly when applied to large datasets, involves extended segmentation sessions and computationally intensive template matching. During these steps, several tables and audio files are generated and managed automatically in the background. To ensure a seamless operation without risks of workflow interruptions and keep reproducibility across analyses, users must adhere to specific prerequisites. First, proper directory management is critical. Working within an R project. Also, we recommend caution when using the ’set_workspace’ over working directories of ongoing projects, as it may add unnecessary files or replace existing data.

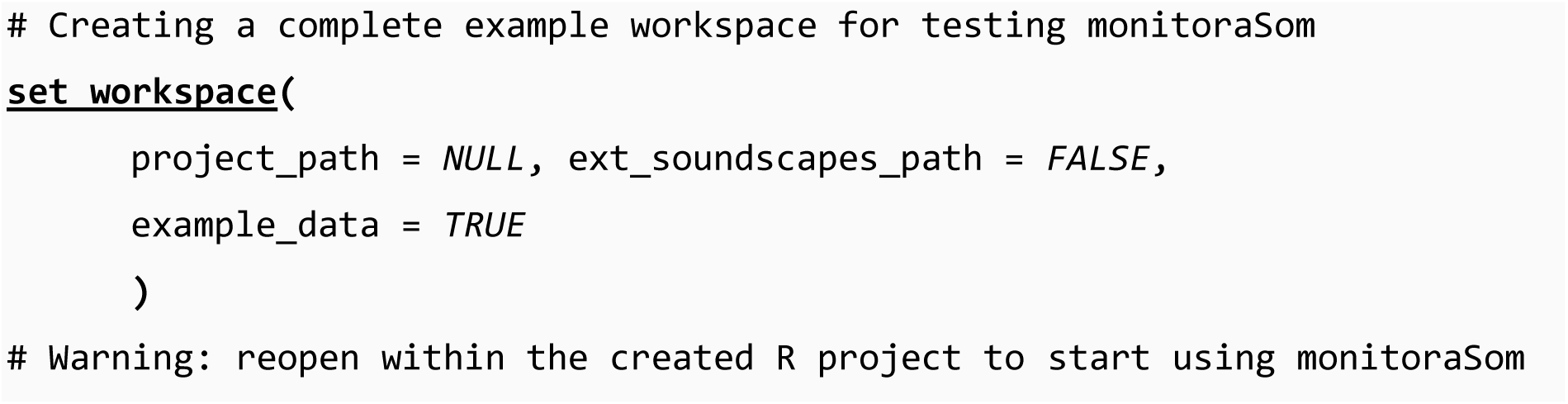

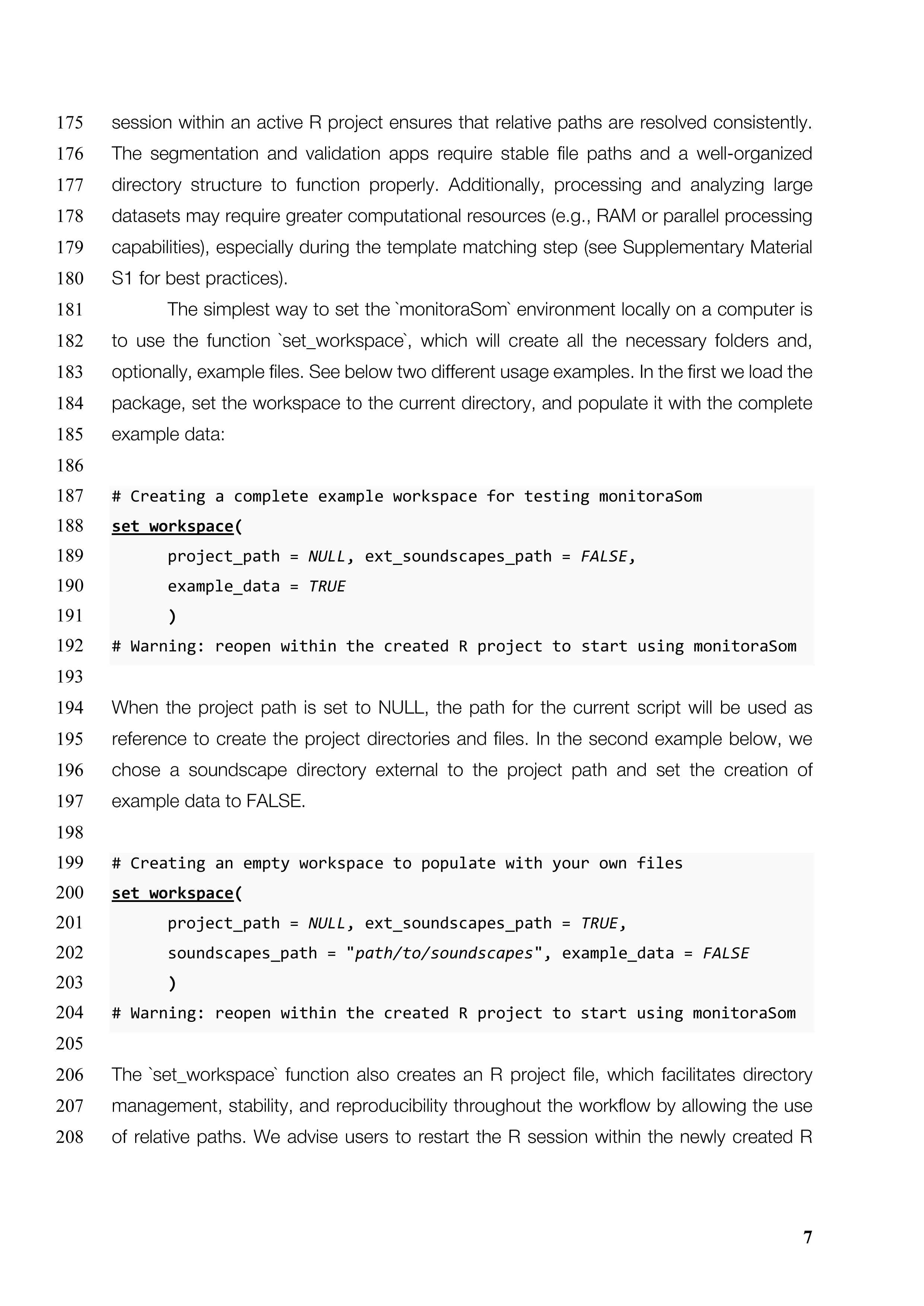

### b. The segmentation and template extraction

The ’launch_segmentation_app’ function initiates an interactive web application implemented with the ’shiny’ package (v.1.10.0; Chang et al. 2024). This tool is designed to facilitate precise annotation of acoustic events by enabling users to manually delimit and label regions of interest (ROls) directly within a spectrogram interface. The user­ friendly and responsive interface was designed to facilitate the process by letting experts focus and dedicate more time into segmentation and less in manual file management and typing. The segmentation data, referred here also as ROI tables and stored as CSV files, has many uses downstream the analysis, but can also be readily used as biological data. The ROls tables can be used for extracting acoustic features with R packages dedicated to bioacoustic analysis, such as ’seewave’ (Sueur et al. 2008) and ’warbleR’ (Araya-Salas & Smith-Vidaurre 2017), with minimal editing. Also, ROI tables can be processed to estimate species richness through accumulation curves, characterizing community composition, or further functional and phylogenetic analyses.

To launch the app, users must define three core parameters: ’project_path’ (directory for output files), ’user’ (observer identifier), and ’soundscapes_path’ (directory containing the target recordings). These parameters can be specified directly in the function call or entered within the app interface, if the app session is already active. When executed in RStudio, the function opens a pop-up window with the app, while users running it in other environments can access the interface via a browser, using the link provided in the R console. The app resolves the paths relative to the currently active R session, ideally within an active R project (see the ’set_workspace’ function).

The app can be launched with minimal setup by using the default settings as shown below, if there are WAV files on the ’soundscapes_path’ and ’cuts_path’ is accessible. From now on we will use the default path of the directories from the example data (see the ’set_workspace’ function for details on how to build it).

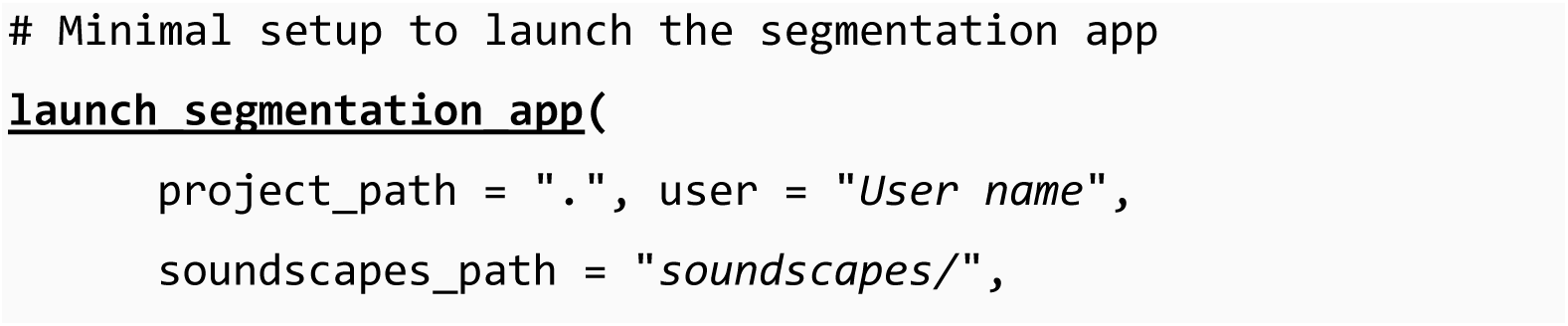
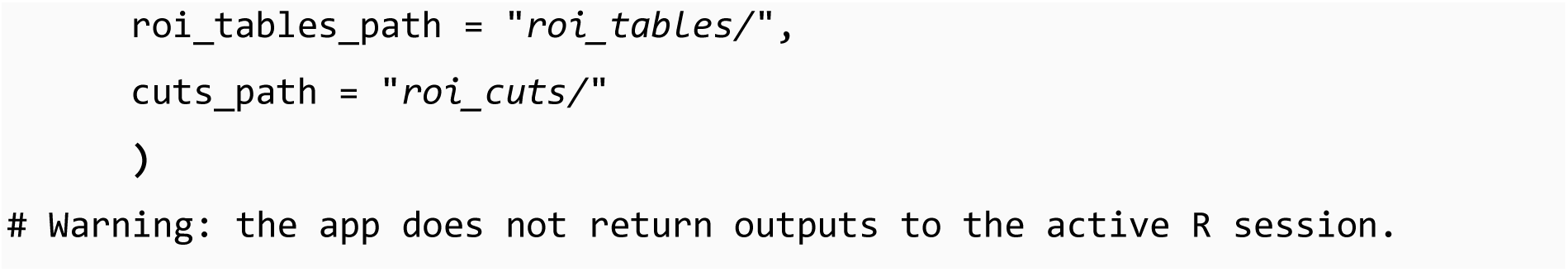

The interface is composed of two parts: a foldable sidebar for session settings (Figure 2a and 2b, black panel on the left) and the main body where the spectrogram and ROI input panels are displayed (Figure 2c and 2d). This layout provides an organized workspace, allowing users to adjust settings while keeping a clear view of the spectrogram and annotation tools. If preferred, most settings can be preset as arguments in the launch function to ensure consistency across segmentation sessions (more details available in Supplementary Material S1).

**Figure 2:**
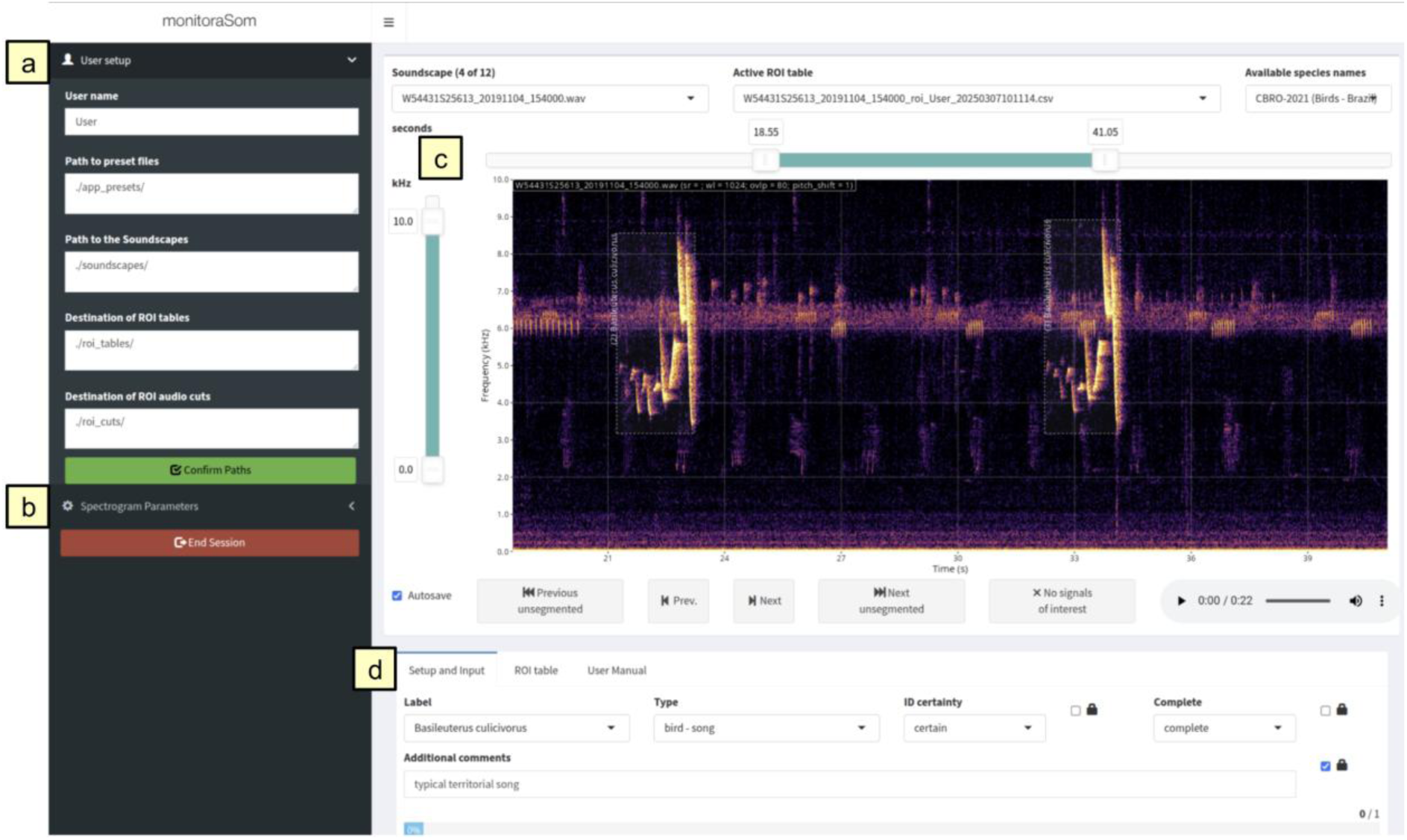
The segmentation app interface is composed of the sidebar that contains the ’User setup’ tab (a) and ’Spectrogram Parameters’ tab (b; see Figure 3 for more details) and the app body that contains the spectrogram panel (c) and the input panel (d).

The sidebar contains two tabs: ’User setup’ (Figure 2.a), where users specify the username, paths for presets and soundscapes, and destination paths for ROI tables and audio cuts; and ’Spectrogram Parameters’ (Figure 2.b and Figure 3), where users can adjust spectrogram and visualization settings, as well as app behavior. Spectrogram settings include dynamic range, which sets the visible energy interval in spectrogram color scale and can be adjusted for better contrast between target signals and background noise. Window length and overlap are also available for adjusting the spectral and temporal resolutions of the spectrogram. Customization options include adjusting the ROI label angle and toggling its visibility to avoid cluttering; horizontal and vertical guides can be added when it is critical for species identification; as well as color scales to match the user’s preferences. Users can also control how the soundscape audio will be played, whether using a player embedded in the app interface, directly by the R session in the background or externally with a third-party software of choice. Additional options allow users to normalize and filter the reproduced audio to playback only the visible frequency band, as well as apply a pitch shift, i.e., a slowdown factor measured in octaves that allows the playback of ultrasonic signals in the human hearing range. It is important to note that none of the actions within the segmentation app take effect over the original soundscape file, and window length and overlap are stored as variables of exported ROI tables.

**Figure 3:**
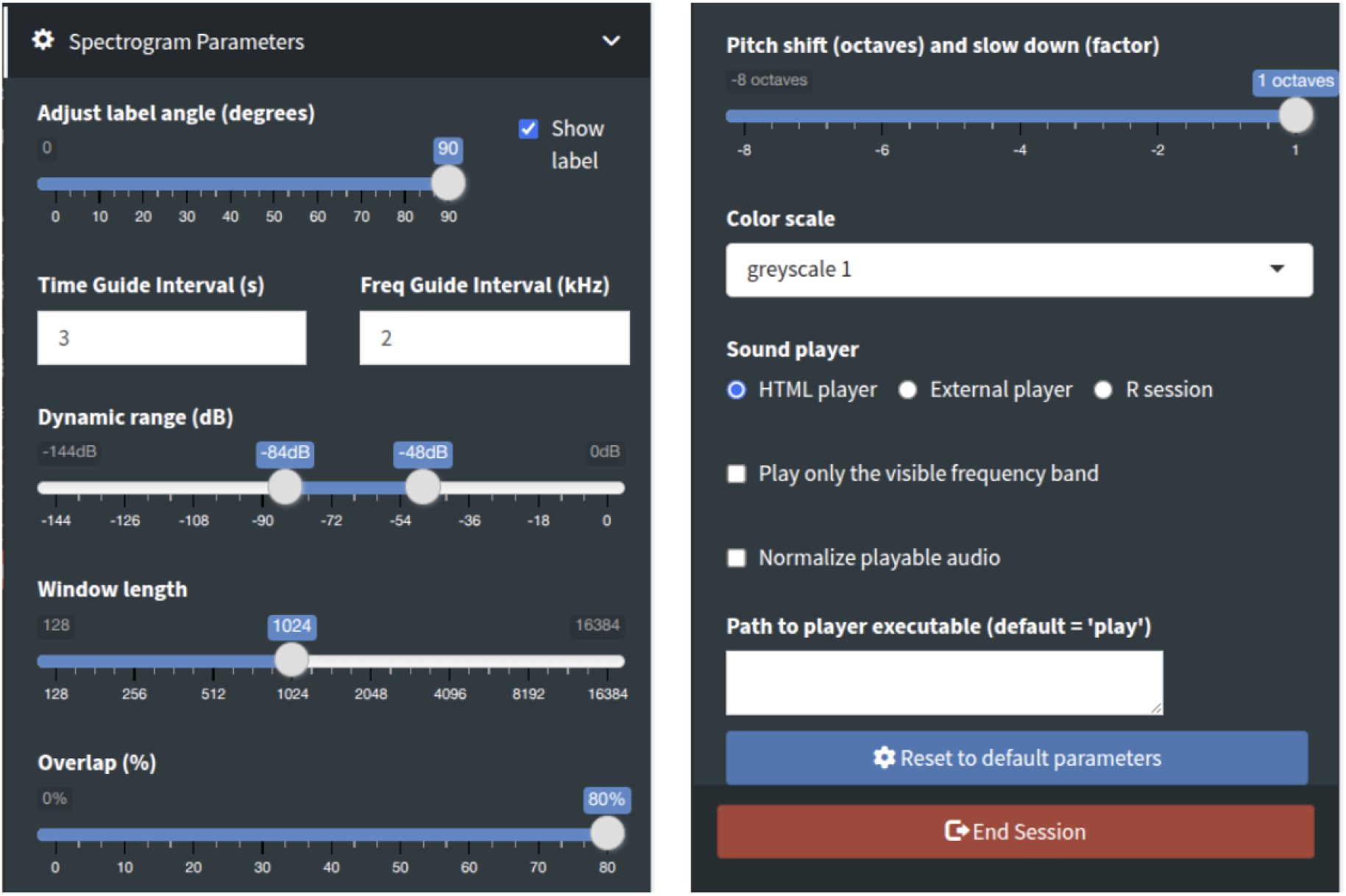
Spectrogram parameters tab at the segmentation app sidebar, showing spectrogram parameters, visualization settings and audio playback settings.

The first panel in the dashboard body (Figure 2.c) shows the interactive soundscape spectrogram in which users can zoom, navigate between recordings, and annotate events using intuitive controls. Immediately below, the second panel contains the input fields for the data that is input on ROls that are about to be made (Figure 2.d). ROls are created by drawing bounding boxes over acoustic signals and storing them with the values provided in the Input tab, which include label, signal type, identification certainty, and signal completeness. All fields support both dropdown selection and direct text input, accepting partial terms and dynamically filtering the available options. Additionally, there is an "Additional comments" field available for users to input any extra information if needed. If the ROI is successfully stored, a persistent box is rendered in the spectrogram and an entry is made in the second tab, called ’ROI table’. Along with each stored ROI, the visualization settings are also stored, enabling researchers to revisit and compare segmentation under the exact conditions in which they were originally defined. In the ’ROI table’ tab, users can edit each ROI and export the current table. More details about hotkeys, buttons and their functionalities are available in the Supplementary Material S1.

Users do not need to manually create a file to store the created ROI - the system generates it automatically. The resulting CSV file name follows a structure format: *"<SOUNDSCAPE name>_roi_<USEMAME> _<TIMESTAMP>.wav".* This ensures consistency and traceability across segmentation sessions. Users can access the available species lists within the app. The default list is "CBRO 2021 - birds Brazil" (Pacheco et al. 2021), but users can load a custom species list or select other lists available by default in ’monitoraSom’. Once a list is chosen, the species will become available in the "Label" dropdown field below the spectrogram. Species’ names will be found by partial matching, allowing the users to quickly find and select the correct ROI label. To add a customized list to the app, users can open the "app_presets/sp_labels.xlsx" file, name the list in the header of the new column, fill this column with the values that must be passed to the ’Label’ dropdown field, save the file, and reload the app for the change to take effect.

The segmentation app is also an essential tool for crafting templates, which in this workflow are derived from segmentation of soundscapes or any other type of recordings containing high quality utterances of the target signal. The first method available for crafting templates is by manually selecting the desired ROls in the ROI table and pressing the button "Export audio of selected ROI" immediately above it. Each template will be exported as a WAV file to the path informed in the ’cuts_path’ with a specific name code

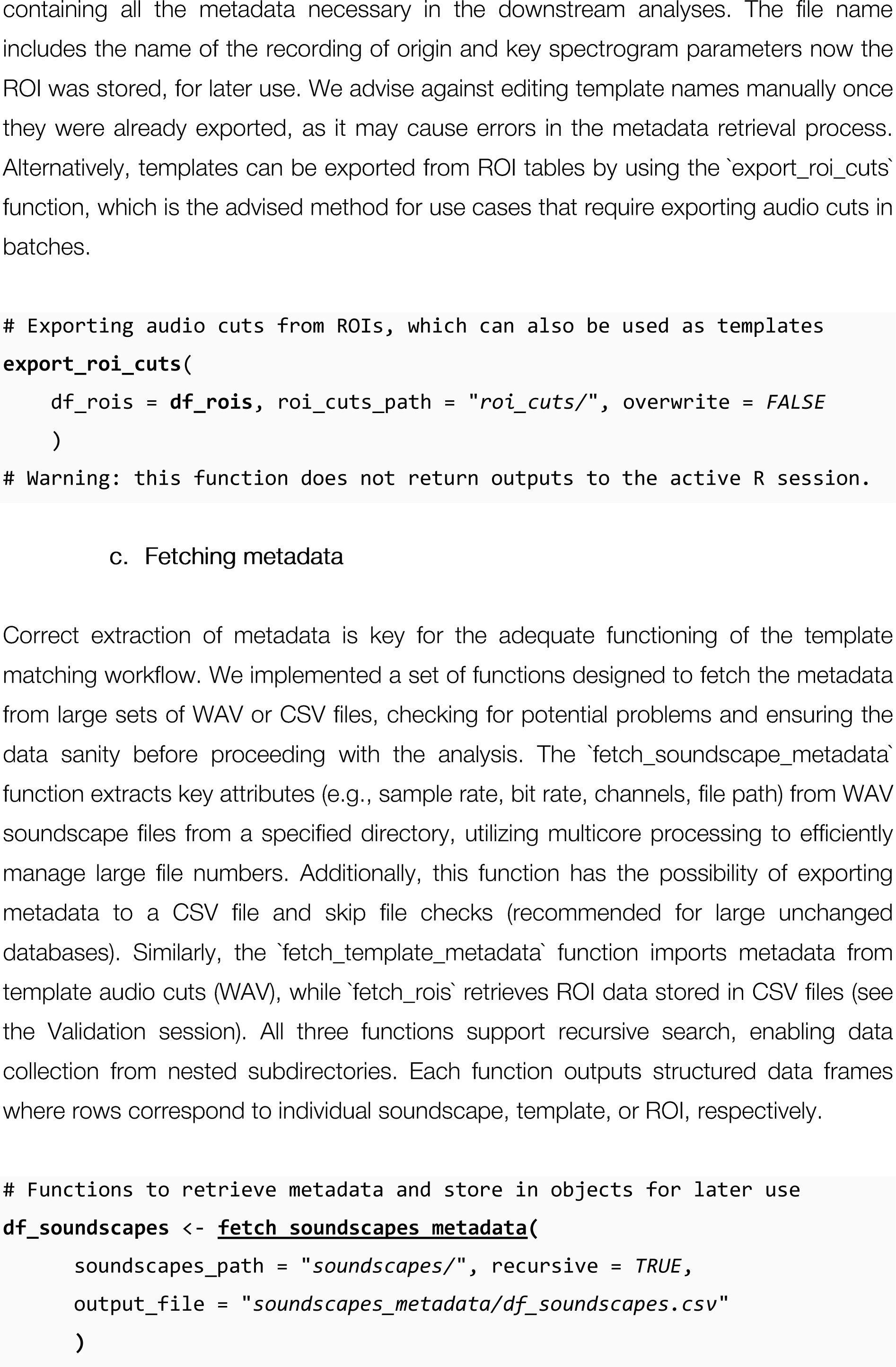
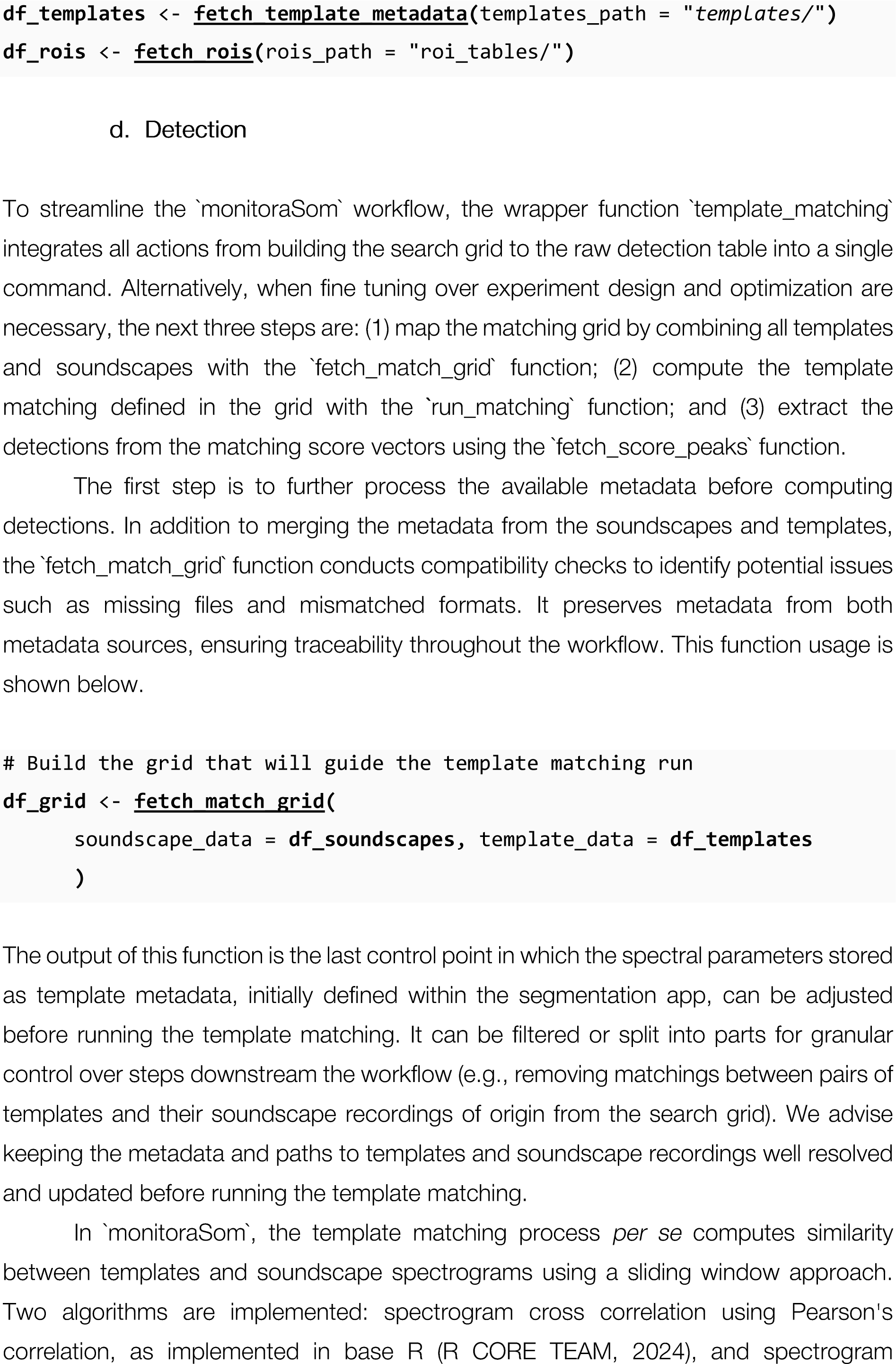
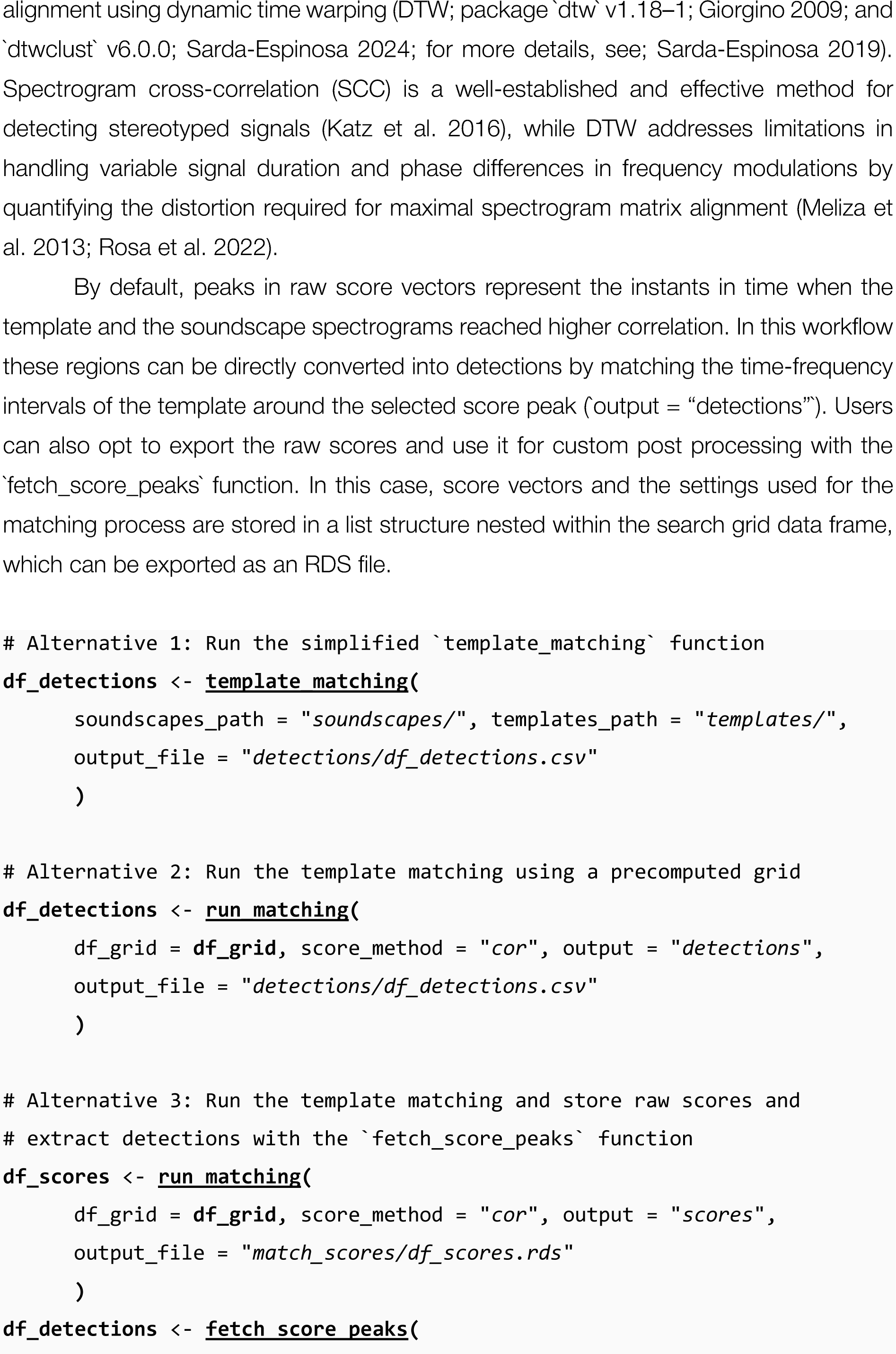
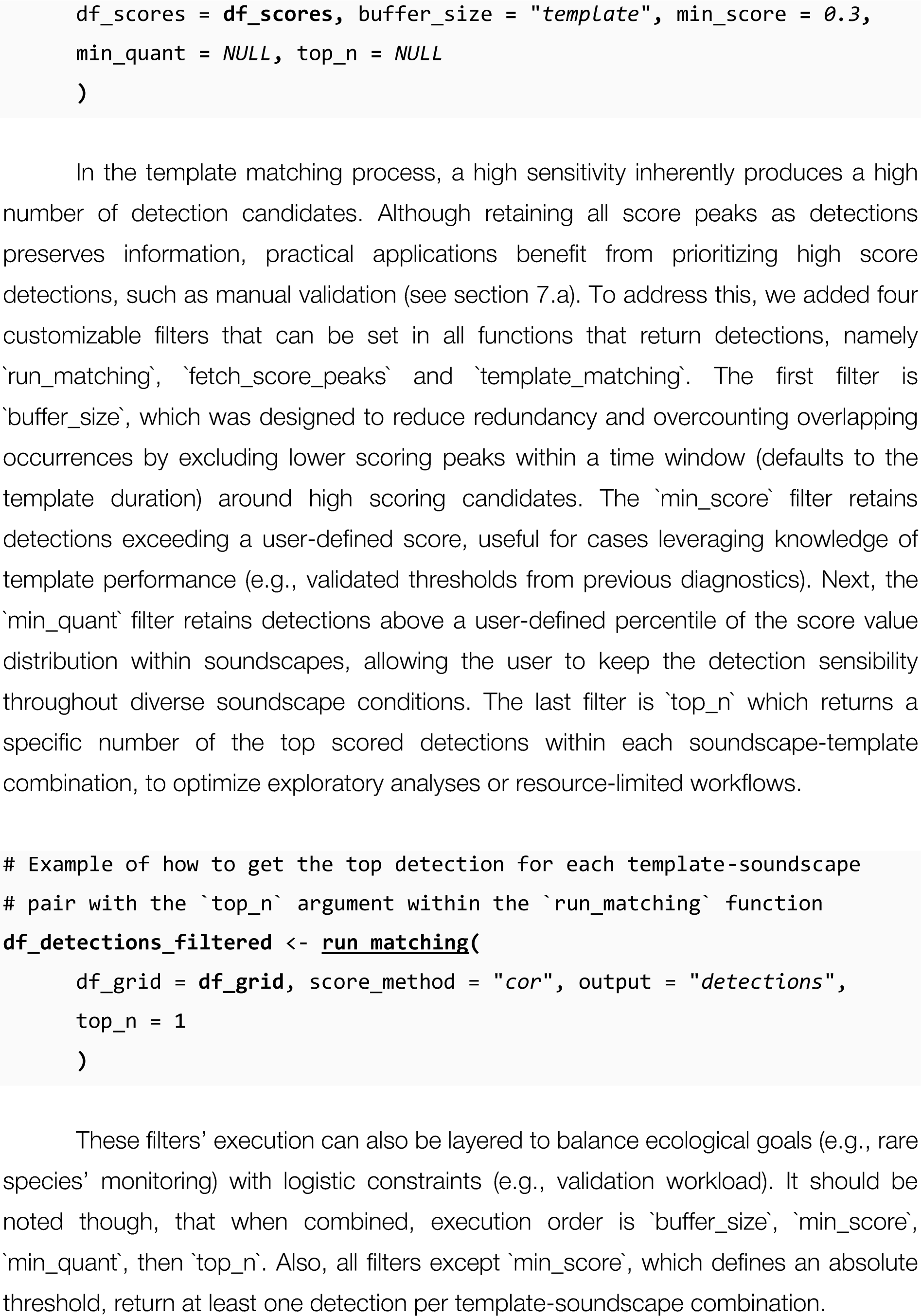
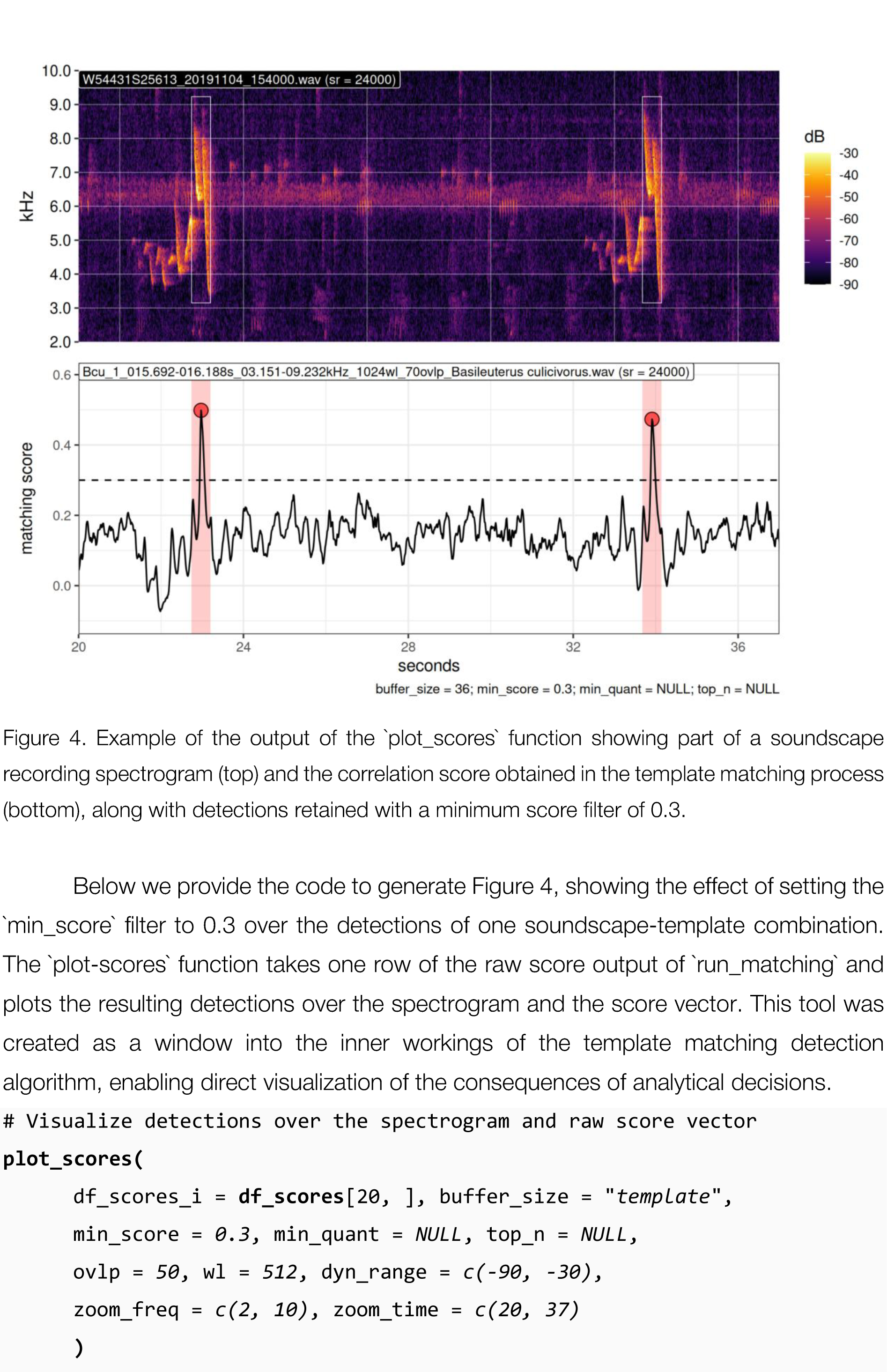
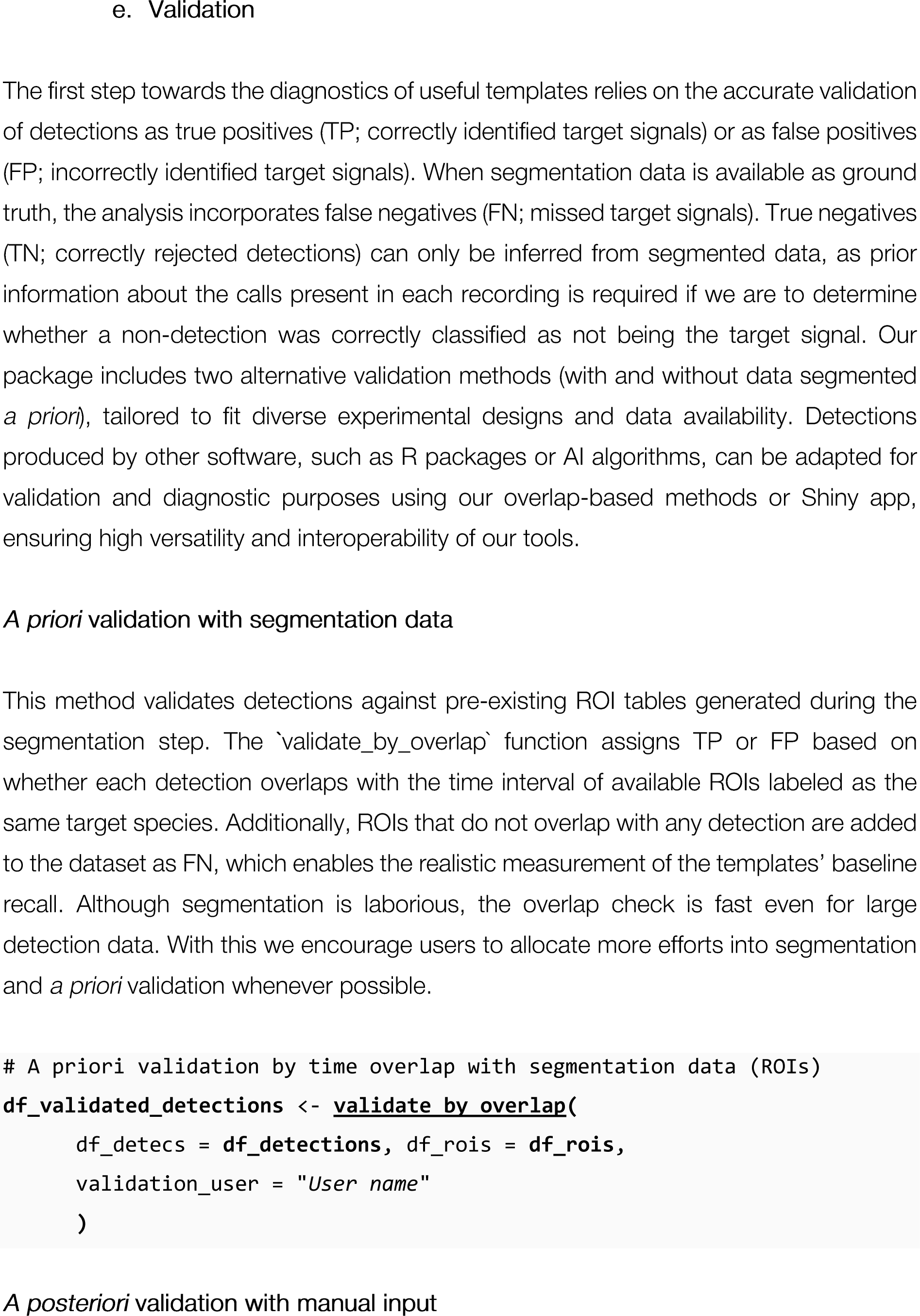
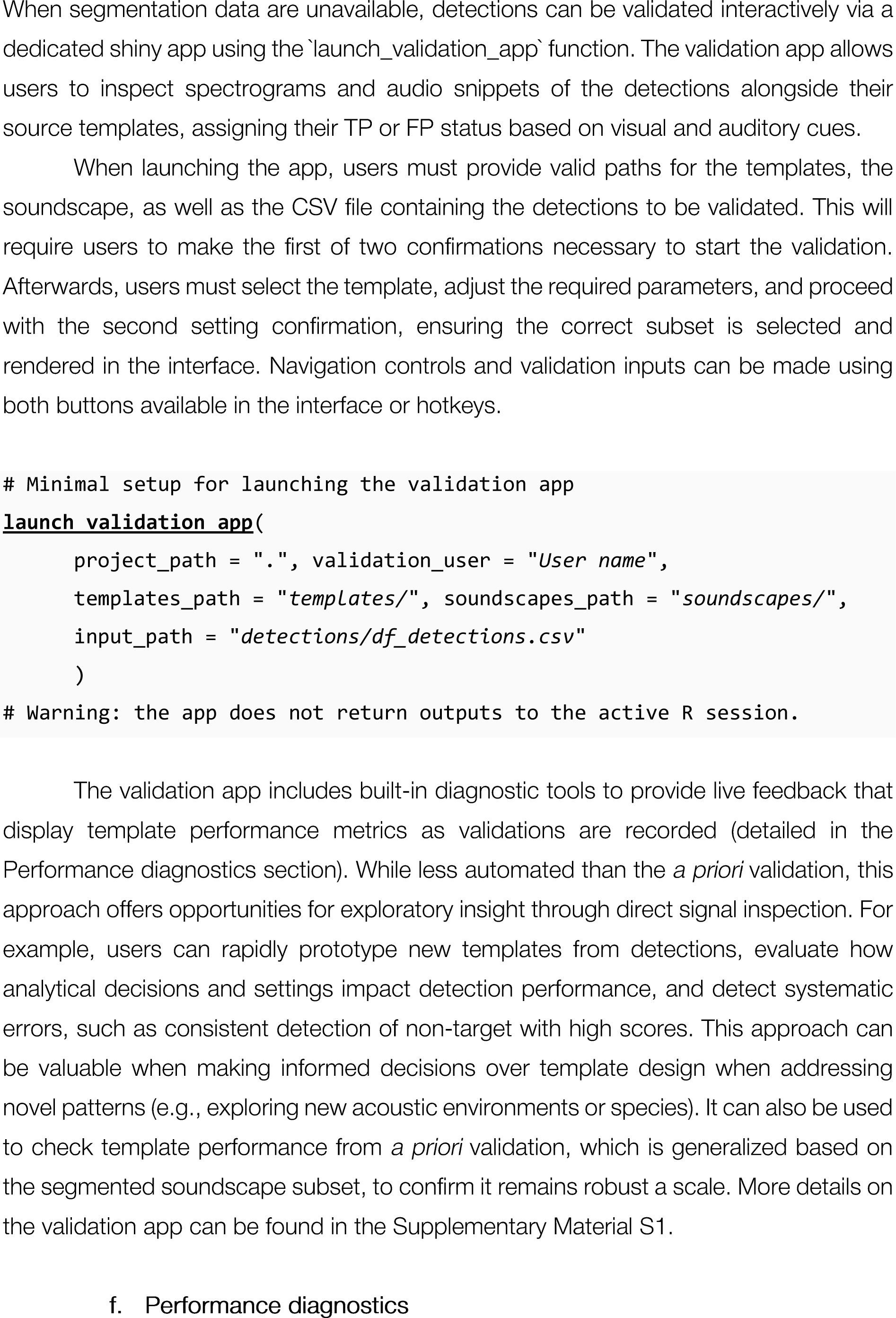
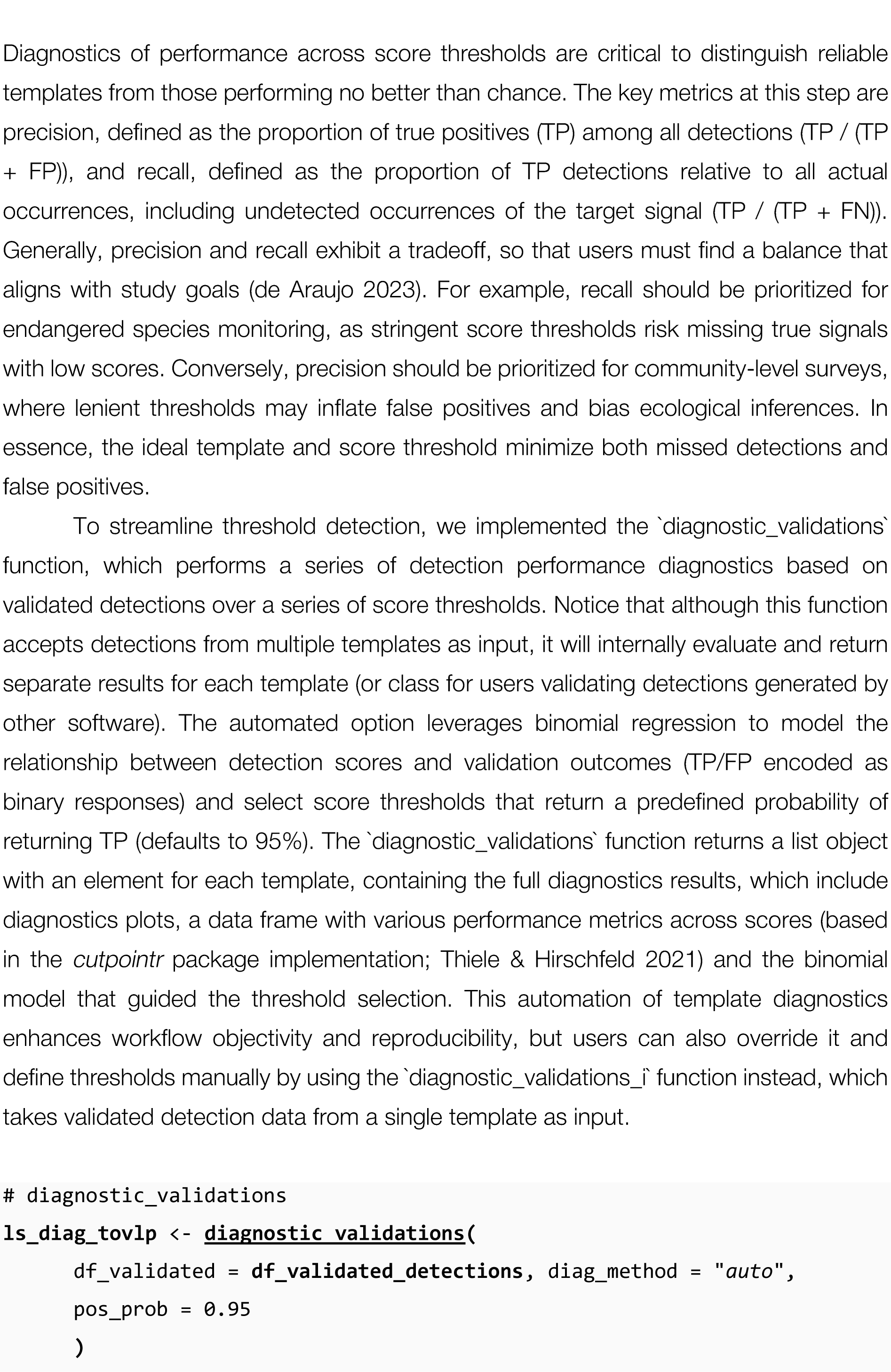
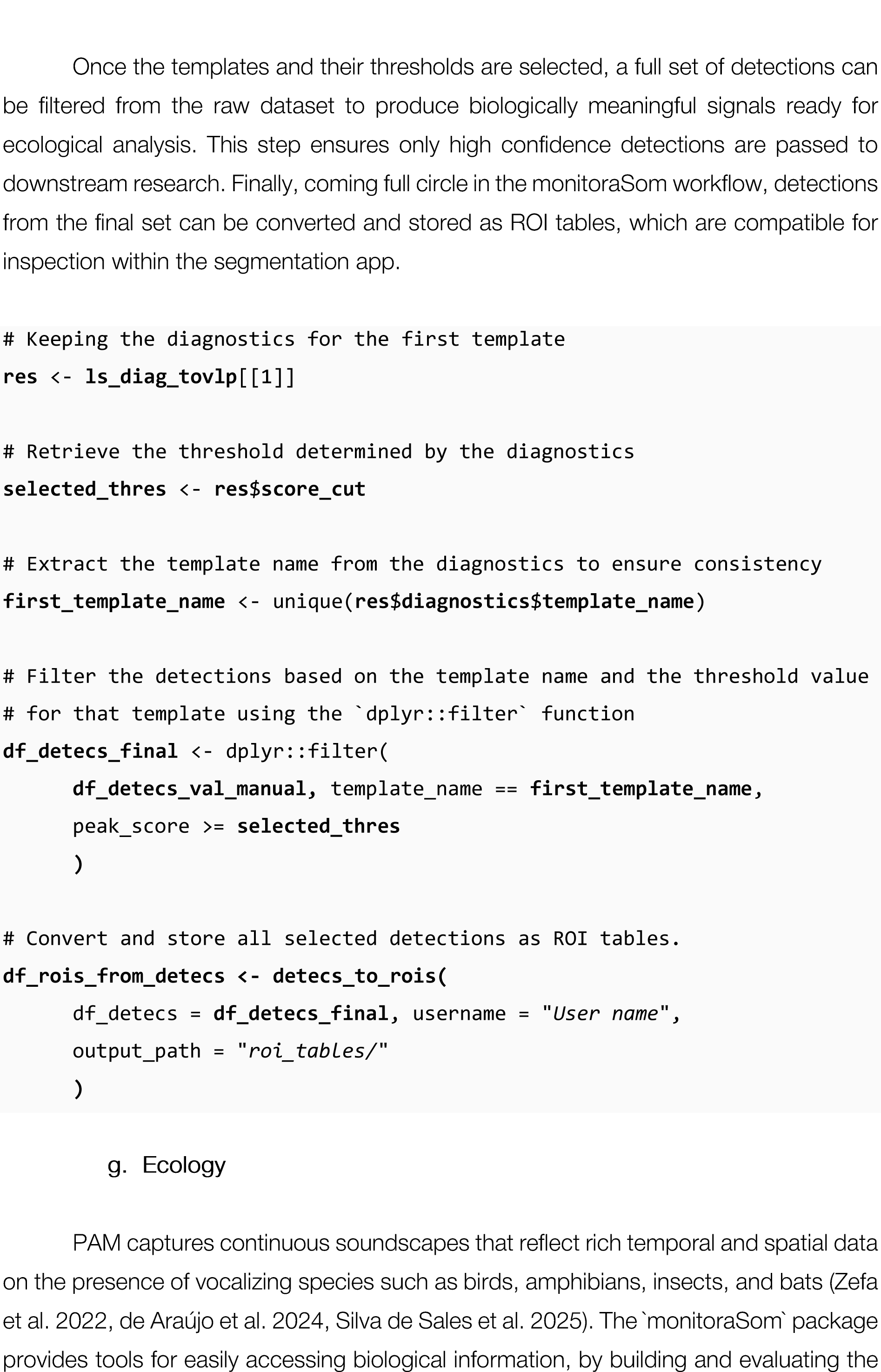

efficiency of automated detectors using metrics such as precision and recall and thereby facilitating the large-scale extraction of biological information from sound recordings. The optimized detection of specific acoustic signals provides valuable evidence for tracking target species over time and space in different ecological scales. A set of detections can be used to test differences on the richness of composition of a community, but also to describe daily and seasonal patterns of specific populations (Perez-Granados & Traba 2021), the presence of rare and cryptic species (de Araujo et al. 2023), and habitat preference assessments (Gangenova et al. 2025, dos Anjos et al. 2022). In fact, raw detection data integrate seamlessly with occupancy models (Campos-Cerqueira & Aide 2016), GLMs, or population dynamics studies.

In an era of accelerating habitat degradation and biodiversity loss, the core aim of ’monitoraSom’ is to facilitate the integration between PAM to test hypotheses both in community and population ecology. By making it broadly accessible and by accelerating the pace of the translation of massive amounts of soundscape data into ecological information, we want to empower researchers to respond to urgent ecological crises.

## 4. DISCUSSION

The tools presented here unify segmentation, detection, validation, and diagnostics into a cohesive, user-friendly workflow. The segmentation and validation apps are designed to minimize errors through controlled interfaces, optimizing efficiency across wide scenarios. Modular workflow design enables comprehensive customization to support exploratory analysis and ensures reproducibility in complex use cases. Automated metadata tracking, standardized file structures, and robust automation of data management, also ensures transparency and scalability that is critical for handling growing volumes of acoustic data and extracting ecological information. We hope ’monitoraSom’ can make the PAM workflow more easily available to newcomers while keeping the analytical rigor, to meet the demands of advanced users. All tools available in ’monitoraSom’ were refined based on extensive feedback from beta testers and field researchers, ensuring that the tool addresses practical challenges encountered while working with diverse taxonomic groups and analytical objectives. We open this software and its workflow for refinement by the user community, expanding its functionalities in favor of improving biodiversity monitoring.

## SOFTWARE AVAILABILITY

The ’monitoraSom’ public repository is available at: https://github.com/ConservaSom/monitoraSom.

## FUNDING

GLMR was funded with postdoctoral scholarship from NAPI Biodiversidade: Servigos Ecossistemicos (Fundagao Araucaria convenio 151/2023), and CBA with postdoctoral fellowship from Consejo Nacional de lnvestigaciones Cientrficas y Tecnicas (CONICET).

## AUTHOR CONTRIBUTIONS

GLMR, implemented the software functions and apps, as well led manuscript writing; GLMR, CBA, and ZJP, conceived the software workflow; CBA, IMDT, CRS, and ZJP contributed significantly with the design and testing of the segmentation and validation apps; LA coordinated the funding from NAPI Biodiversidade for this research. All the authors contributed to writing this manuscript.

## Notes

https://github.com/ConservaSom/monitoraSom

### Competing Interest Statement

The authors have declared no competing interest.

