## Supplementary Material S1 for "monitoraSom: an easy path from Soundscapes to Ecology"

### monitoraSom

Rosa, G. L. M.; Zurano, J. P.; Torres, I. M. D.; Simões, C. R.; dos Anjos, L.; de Araújo, C. B.

2025-06-23

#### Overview

Here we provide an overview of the R package **monitoraSom**. The package is designed to facilitate passive acoustic monitoring (PAM) and bioacoustic analyses by providing tools for processing, analyzing, and visualizing sounds for ecological studies. It includes functions for efficiently segmenting audio recordings, extracting acoustic features, building spectrograms and waveforms, and performing batch processing of large datasets. These capabilities make monitoraSom particularly useful for sound ecologists.

In this vignette, we present a complete workflow for template matching analysis, demonstrating how to detect specific acoustic signals within a set of soundscape recordings. The workflow begins with structuring a working directory locally to ensure that all files are easily accessible. We use high-fidelity recordings of *Basileuterus culicivorus* to create acoustic templates that will serve as reference sounds in the template matching analysis. The template matching builds a cross-correlation analysis to search for moments in which the sounds in a set of soundscapes are similar to that of a template. The validation *a posteriori* helps establish a minimum correlation threshold that will most likely contain a target vocalization. The detections validated as true occurrences (true positives) can then be exported for ecological analysis, such as occupancy models or vocal diel cycle descriptions. The workflow described in this vignette provides a tool for population-level bioacoustic studies.

#### Setting R environment

##### *Working directory structure*

Once the `monitoraSom` package is installed, you can load the libraries into your R environment. In this vignette, we use the packages `monitoraSom` and `dplyr`. After executing the `library()` function, the tools from `monitoraSom` and `dplyr` will be available for use in your R session. These tools enable effective handling of acoustic monitoring data and data wrangling tasks. The packages can be loaded with the following command:

```
library(monitoraSom)
library(dplyr)
library(patchwork)
```

##### *Folder structure*

monitoraSom is designed to handle large-scale passive acoustic data, where files must be systematically stored for easy access and to avoid problems during analysis. The files are organized within a working directory, so our first task is to set a local folder as the working directory. To examine your current working directory, use the following command:

```
current_dir <- getwd()
current_dir
```

```
## [1] "/home/grosa/R_repos/monitoraSom/example/vignette_01_Essentials"
```

If you are running this script from an R Markdown file in the RStudio IDE, the current working directory is the folder containing the R Markdown file. However, if you need to change it, you can use the `setwd()` function directly.

```
setwd("path/to/your/working/directory")
```

Alternatively, you can use the following command to get the directory of the current file and set it as the working directory:

```
current_dir <- dirname(rstudioapi::getSourceEditorContext()$path)
setwd(current_dir)
```

Within the working directory, monitoraSom uses a folder structure that handles multiple file types, from sound files to data tables. While you can set all directories manually, the easiest way to establish monitoraSom's directory structure is by using the `set_workspace()` function. This function automatically creates all necessary directories using a standard structure. If you choose to include the demo data as used here, you can set the parameter `example_data = TRUE`. This will copy all essential files from this demo into the working directory. Note that some directories are not required in this example and can be set to NULL.

```
set_workspace(
  project_path = current_dir, example_data = TRUE,
  app_presets_path = "./app_presets/",
  soundscapes_path = "./soundscapes/",
  soundscapes_metadata_path = NA,
  recordings_path = "./recordings/",
  roi_tables_path = "./roi_tables/",
  roi_cuts_path = NA,
  templates_path = "./templates/",
  templates_metadata_path = NA,
  match_grid_metadata_path = NA,
  match_scores_path = NA,
  detections_path = "./detections/",
  detection_cuts_path = NA,
  detection_spectrograms_path = NA,
  validation_outputs_path = "./validation_outputs/",
  validation_diagnostics_path = "./validation_diagnostics/"
)
```

### Launching the segmentation app

The segmentation app is a multipurpose tool that can be used for various objectives. It serves as a quick visualization tool to inspect recordings, list species present in sound files, evaluate species repertoire, and measure specific behavioral aspects of calls.

Now that the directories are in place, let's launch the segmentation app with a minimal setup. The essential parameters for launching the app are the paths to the project folder (`project_path`), the path to the soundscapes folder (`soundscapes_path`), the path where to store the ROI tables folder (`roi_tables_path`), and the user identification (in this case set as `user`). You can note your ideal settings and establish the paths directly in the `launch_segmentation_app()`. The app will be launched by executing the command below.

```
launch_segmentation_app(
    project_path = ".", user = "User Name",
    soundscapes_path = "./soundscapes/",
    roi_tables_path = "./roi_tables/"
)
```

Note that once the app is open, it will be rendered in a pop-up window. Check if the window dimensions are appropriate for your screen size. If not, you can resize the window by dragging its edges. If the window is too small or the screen resolution is too low, the app interface elements might not display properly. If the app is rendered correctly, make sure to check the paths on the sidebar before pressing the 'confirm path' button (red in Figure 1) to render the spectrogram of the first file. For your convenience, the sidebar can be hidden, providing a larger workspace for ROI selection.

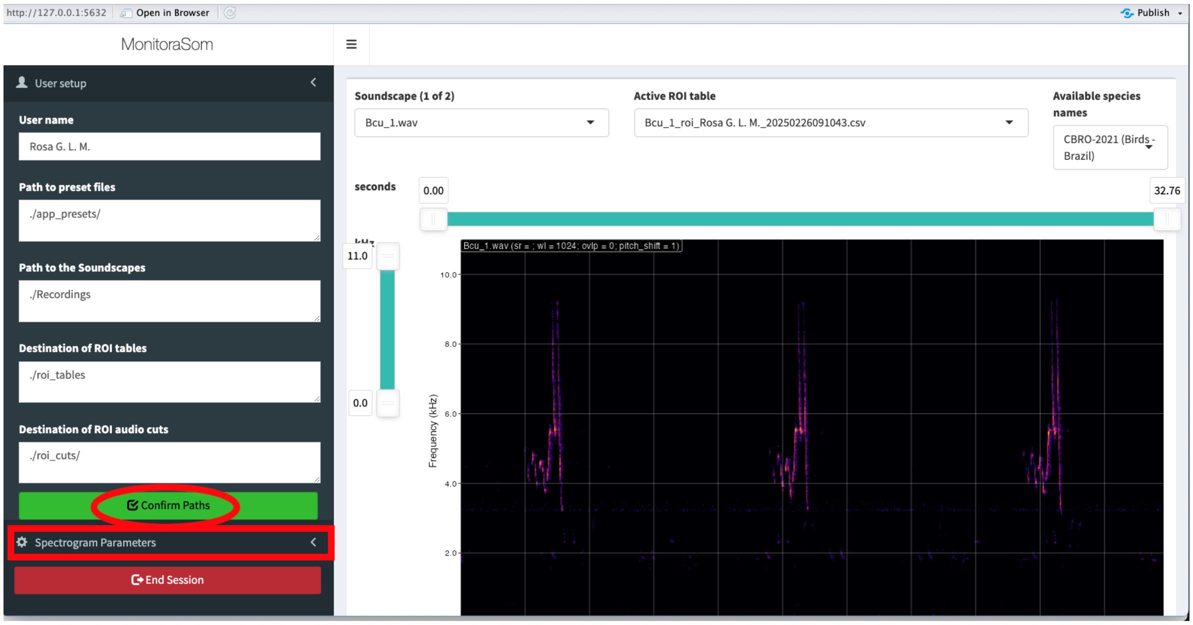


Figure 1: The segmentation app highlighting the confirmation button, necessary to build the first spectrogram, and the spectrogram parameters menu.

In the spectrogram parameters menu (highlighted with a red rectangle in Figure 1), you can access spectrogram parameters to adjust the display of the spectrogram. These parameters can be controlled within the app for immediate use or preset in the function arguments before launching the app. Within the menu, you can also reset to values passed to the launch function using the "Reset to default parameters" button, located at the bottom of the spectrogram settings tab in the sidebar (Figure 2). Remember to note your preferred settings before closing the app to pass them as new defaults to the launch function.

Now that spectrogram parameters are set, you can visually search for a signal of interest in the spectrogram, use play button (spacebar) to listen to it, and select a region of interest (ROI) using the mouse. Once you press 'E' on the keyboard, the selection's metadata will be stored as one row on the ROI table of the current soundscape. Several parameters of the ROI are stored at this point, including the minimum and maximum frequency, as well as its start and end within the soundscape recording. However, there are other data that can be added to the ROI before it is stored. These data include several relevant pieces of information on the label of the ROI, such as the species, call type, and the certainty of the label. The input fields to add this information to a ROI are available in a panel below the spectrogram. You select the species label from a specific list of taxa, which can be found in the `"/app_presets"` folder, as an XLSX file. Out of the box, `monitoraSom` offers multiple species lists from multiple authorities, stored as columns in the XLSX file. Our example includes the list of Brazilian birds from CBRO (2021) and Argentinean birds by Aves Argentinas. Other lists are also available, such as the list of amphibians from the Brazilian Herpetological Society (2021) and the bats of Brazil from CLMB (2020). Among the inputs for the current ROI, you can also select the signal type from a wide range of preloaded repertoire components, from a bat feeding buzz to a frog advertisement call or a bird song-duet. Ensure that the metadata is as complete as possible, including information on label certainty, which is crucial for quality control. Misidentifications might be difficult to spot at other steps of the workflow. Therefore, entries with uncertain identification should be clearly identified as such, allowing specialists to review them as needed.

In this example, we preloaded the ROI tables with all songs of *B. culicivorus* present in the soundscape recordings. It is good practice to identify and label all occurrences of the target species. The songs of *B. culicivorus*, like many animal vocalizations, are structured across multiple hierarchical levels. For instance, birds can produce groups of notes (units of acoustic events) that together form song phrases. A bird recording that captures repeated phrases contains the highest level of vocal organization and can be considered a 'complete' recording. Therefore, the hierarchical level at which the segmentation is performed depends on the objective of the analysis. In this example, we will segment the complete songs of *B. culicivorus* into substructures A, B, and C, which are consistent across multiple songs.

Let's launch the app again with a more complete setup. Instead of segmenting soundscape recordings, we will now segment focal recordings from the target species. Observe the structure of the song of *B. culicivorus* in the spectrogram and how consistent the substructures are across multiple songs. Avoid excluding the preloaded ROIs from this example to ensure the consistency of the next steps.

```r
launch_segmentation_app(
  project_path = ".", user = "User Name",
  soundscapes_path = "./recordings/",
  roi_tables_path = "./roi_tables/",
  time_guide_interval = 10, freq_guide_interval = 1,
  dyn_range = c(-60, -24), wl = 1024, ovlp = 70,
  zoom_freq = c(2.9, 10.1), color_scale = "greyscale 1"
)
```

Let's review how the segmentation process works:

1. *Launching the app.* Launch the app with the `launch_segmentation_app()` function, passing the correct paths to the `project_path`, `soundscapes_path`, and `roi_tables_path` arguments.
2. *Confirm paths.* Click the "Confirm paths" button. If the paths are correct, the spectrogram of the first soundscape will be displayed.
3. *Navigating files.* Zoom in the frequency range using the W and S keys, or by clicking and dragging the time and frequency sliders until the desired range is selected. The visible frequency band of the spectrogram affects data retrieval, so it is important to be consistent throughout the analysis. Proceed if you are satisfied with the spectrogram window settings.
4. *Searching for signals.* Search for a signal of interest in the spectrogram and fill in the metadata of the ROI you are about to create as completely as possible.
5. *Drawing a ROI.* Click and drag the mouse to draw a rectangular selection around the signal of interest in the spectrogram. Press the `E` key to store this selection as a ROI in the current ROI table. The metadata fields you previously filled will be automatically applied to any new ROIs you create, so you don't need to re-enter this information for each selection unless you want to change some of the information.

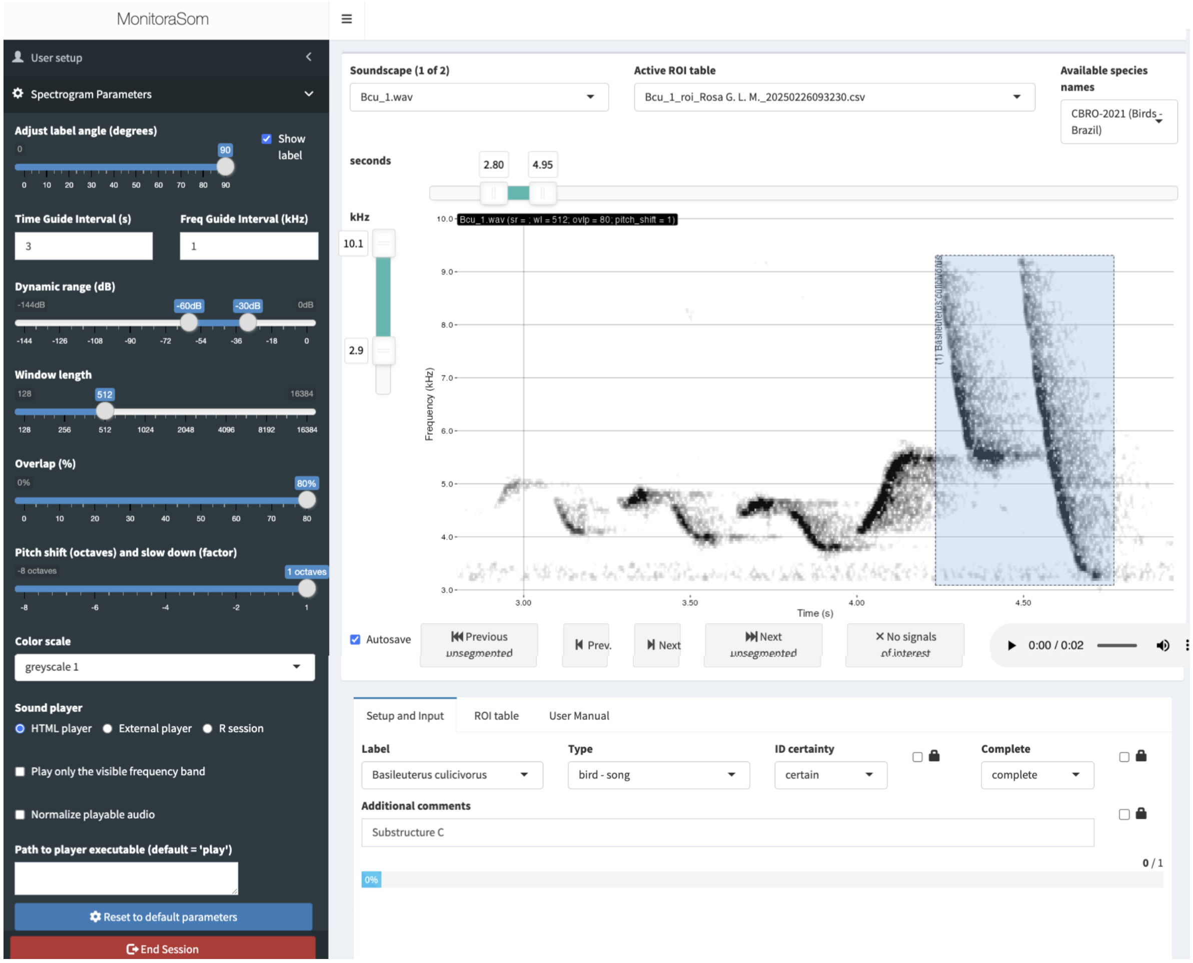


Figure 2: The segmentation app highlighting the selection of the structure ‘C’ of *B. culicivorus* song.

6. *Follow the workflow.* Repeat steps 3 and 4 for each signal of interest in the spectrogram until the objective of the segmentation is achieved (all ROIs from the current soundscape can be reviewed in the ROI table tab below the spectrogram). This process is designed to be highly efficient if you segment signals from one species at a time.
7. *Navigate soundscapes.* Navigate between the recordings using the `Z` and `C` keys, repeating the steps above until all the soundscapes are segmented. If the autosave option is enabled, the ROI tables will be saved automatically as CSV files in the path passed to the `roi_tables_path` argument. Otherwise, you will need to save manually before changing files to avoid data loss (ROI data is not stored within the R session). Navigation between soundscapes is also possible using the soundscape dropdown menu at the top of the spectrogram.

### Building templates

#### *Manual template extraction*

In cases such as this, with multiple songs of the target species, you can use R to extract basic parameters for further analysis, or you can use the segmentation app to extract templates interactively. We will first demonstrate the method to extract templates manually from within the segmentation app using the 'export audio files' button (see the figure below), but there is also a dedicated function (more suited for large-scale use cases) that we will demonstrate later.

In our case, multiple substructures of the song of *B. culicivorus* were selected for use as templates. Visualizing the spectrogram and ROIs within the app helps in selecting the right substructures and performing more direct quality control over the template extraction process. Once the ROIs are selected in the ROI table tab, you can export all the ROIs as templates at once for later use.

Check the `roi_cuts` folder to see the templates extracted from the soundscape recordings.

#### *Automatic audio extraction*

Although we use a small set of soundscape recordings in this example, the segmentation process for template extraction demonstrated is the same as the one used for large-scale template matching analysis. Let's see how to do it using the `export_roi_cuts()` function. First, we need to import the ROI tables back into R. The `fetch_rois` function will return a dataframe that aggregates all the ROIs in the `roi_tables` directory.

```
df_rois <- fetch_rois(rois_path = "./roi_tables/")
glimpse(df_rois)
```

```
## Rows: 71
## Columns: 19
## $ soundscape_path      <chr> "./recordings/Bcu_1.wav", "./recordings/Bcu_1.wav~
## $ soundscape_file      <chr> "Bcu_1.wav", "Bcu_1.wav", "Bcu_1.wav", "Bcu_1.wav~
## $ roi_path             <chr> "/home/grosa/R_repos/monitoraSom/example/vignette~
## $ roi_file             <chr> "Bcu_1_roi_User_20250226101835.csv", "Bcu_1_roi_U~
## $ roi_user             <chr> "User", "User", "User", "User", "User", "User", "~
## $ roi_input_timestamp  <chr> "2025-02-26 10:19:57", "2025-02-26 10:20:12", "20~
## $ roi_label            <chr> "Basileuterus culicivorus", "Basileuterus culiciv~
## $ roi_start            <dbl> 4.236420, 15.691662, 27.506793, 3.794958, 15.2593~
## $ roi_end              <dbl> 4.741796, 16.188014, 28.016681, 4.189475, 15.6949~
## $ roi_min_freq         <dbl> 3.150896, 3.150896, 3.057099, 3.659274, 3.561569,~
## $ roi_max_freq         <dbl> 9.200792, 9.232058, 9.232058, 5.629660, 5.743649,~
## $ roi_type             <chr> "bird - song", "bird - song", "bird - song", "bir~
```

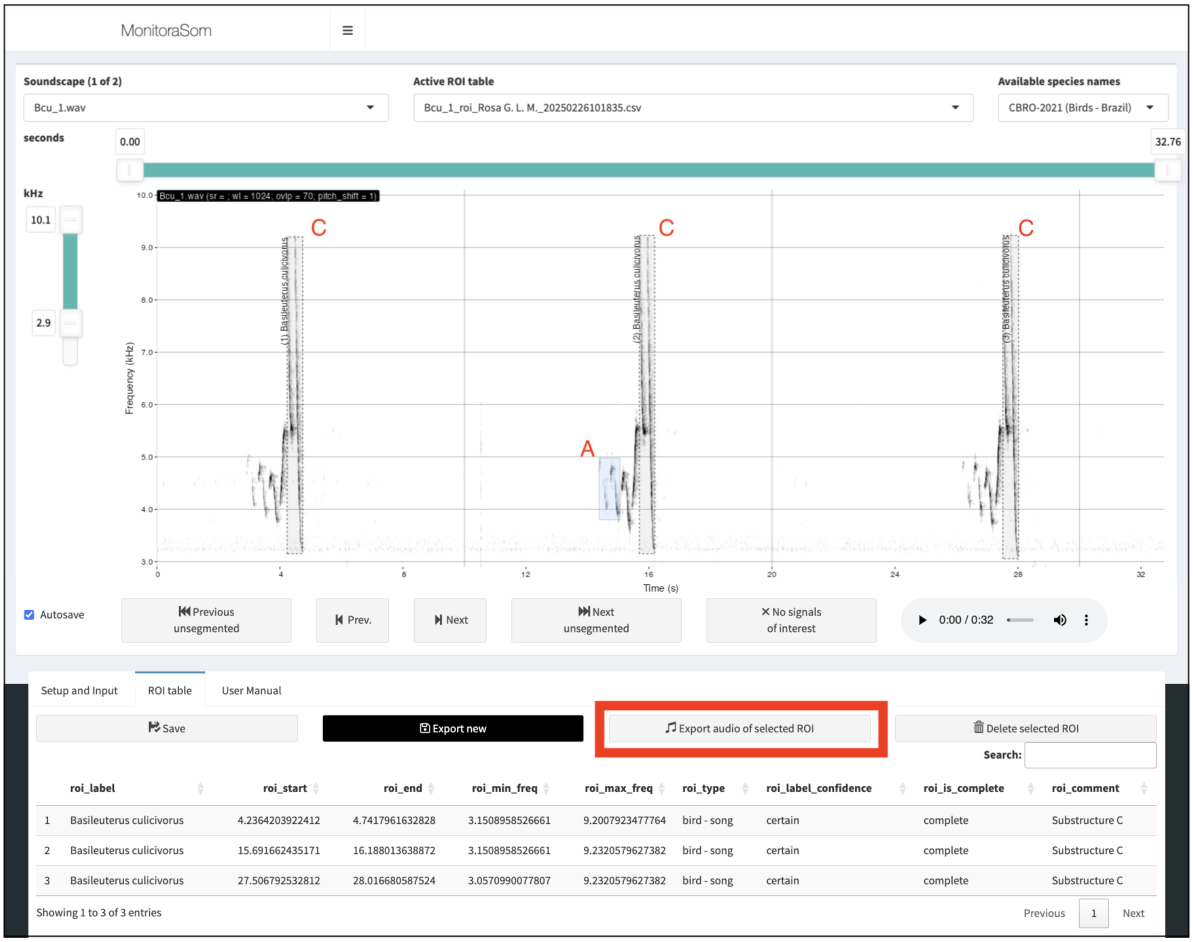


Figure 3: The segmentation app highlighting a complete recording, which includes multiple songs of *B. culicivorus*.

```
## $ roi_label_confidence <chr> "certain", "certain", "certain", "certain", "cert~
## $ roi_is_complete      <chr> "complete", "complete", "complete", "complete", "~
## $ roi_comment          <chr> "Substructure C", "Substructure C", "Substructure~
## $ roi_wl               <int> 1024, 1024, 1024, 1024, 1024, 1024, 1024, 1024, 1~
## $ roi_ovlp             <int> 70, 70, 70, 70, 70, 70, 70, 70, 70, 70, 70, 70, 7~
## $ roi_sample_rate      <int> 24000, 24000, 24000, 24000, 24000, 24000, 24000, ~
## $ roi_pitch_shift      <int> 1, 1, 1, 1, 1, 1, 1, 1, 1, 1, 1, 1, 1, 1, 1, 1, 1~
```

Using the `export_roi_cuts()` function requires a ROI table with the cuts (or templates) that will be exported, which we get using the `fetch_rois()` function. Let's quickly check what recordings are represented in the `df_rois` dataframe.

```
count(df_rois, soundscape_file)
```

```
##                      soundscape_file n
## 1                          Bcu_1.wav 9
## 2                          Bcu_2.wav 9
## 3  W54431S25613_20191102_055000.wav 4
## 4  W54431S25613_20191102_064000.wav 4
## 5  W54431S25613_20191104_154000.wav 4
## 6  W54431S25613_20191105_060000.wav 3
## 7  W54443S25620_20191105_054000.wav 6
## 8  W54448S25622_20191101_073000.wav 8
## 9  W54448S25622_20191101_125000.wav 8
## 10 W54448S25622_20191102_070000.wav 4
## 11 W54448S25622_20191103_055000.wav 7
## 12 W54448S25622_20191104_171000.wav 4
## 13 W54448S25623_20191102_064000.wav 1
```

We can promptly see that it includes both the ROIs from the soundscape recordings and the ROIs from the focal recordings. We have collected three substructures for each song (3), of three different songs (3), from two distinct sound recordings (2), totaling 18 templates. We also have 53 ROIs from the soundscape recordings, which have no information about the substructure in the `roi_comment` column. It is important to note that filtering the desired ROIs is necessary to avoid exporting all ROIs from this dataframe. Let's check the number of ROIs per substructure collected during the segmentation process to see if we can use this information to guide the filtering process.

```
count(df_rois, roi_comment)
```

```
##      roi_comment  n
## 1 Substructure A  6
## 2 Substructure B  6
## 3 Substructure C  6
## 4           <NA> 53
```

Yes, we can easily identify that ROIs from soundscape recordings have no information about the substructure in the `roi_comment` column. We can use this information to filter the ROIs that will be exported as templates. The filtering strategy is open to the user's creativity. In the example below, we filter the ROIs that have the word "Substructure C" in the content of the `roi_comment` column. This will ensure at a single step that ROIs from soundscape recordings are not included, and only the templates of the substructure are exported.

```
df_rois_filtered <- df_rois %>%
    filter(grepl("Substructure C", roi_comment))
glimpse(df_rois_filtered)
```

```
## Rows: 6
## Columns: 19
## $ soundscape_path      <chr> "./recordings/Bcu_1.wav", "./recordings/Bcu_1.wav~
## $ soundscape_file      <chr> "Bcu_1.wav", "Bcu_1.wav", "Bcu_1.wav", "Bcu_2.wav~
## $ roi_path             <chr> "/home/grosa/R_repos/monitoraSom/example/vignette~
## $ roi_file             <chr> "Bcu_1_roi_User_20250226101835.csv", "Bcu_1_roi_U~
## $ roi_user             <chr> "User", "User", "User", "User", "User", "User"
## $ roi_input_timestamp  <chr> "2025-02-26 10:19:57", "2025-02-26 10:20:12", "20~
## $ roi_label            <chr> "Basileuterus culicivorus", "Basileuterus culiciv~
## $ roi_start            <dbl> 4.236420, 15.691662, 27.506793, 4.443569, 14.3930~
## $ roi_end              <dbl> 4.741796, 16.188014, 28.016681, 4.807830, 14.7940~
## $ roi_min_freq         <dbl> 3.150896, 3.150896, 3.057099, 2.900000, 2.975339,~
## $ roi_max_freq         <dbl> 9.200792, 9.232058, 9.232058, 9.763573, 9.554146,~
## $ roi_type             <chr> "bird - song", "bird - song", "bird - song", "bir~
## $ roi_label_confidence <chr> "certain", "certain", "certain", "certain", "cert~
## $ roi_is_complete      <chr> "complete", "complete", "complete", "complete", "~
## $ roi_comment          <chr> "Substructure C", "Substructure C", "Substructure~
## $ roi_wl               <int> 1024, 1024, 1024, 1024, 1024, 1024
## $ roi_ovlp             <int> 70, 70, 70, 70, 70, 70
## $ roi_sample_rate      <int> 24000, 24000, 24000, 24000, 24000, 24000
## $ roi_pitch_shift      <int> 1, 1, 1, 1, 1, 1
```

Now we can export the templates using the `export_roi_cuts()` function. Note that we have routed the output to the `./templates` directory.

```
export_roi_cuts(
    df_rois = df_rois_filtered,
    roi_cuts_path = "./templates",
    overwrite = TRUE
)
```

Although templates can be created manually from within the segmentation app, using the `export_roi_cuts()` is the most straightforward way to export the templates. The file name contains all the necessary information for a standalone template, making it easily used downstream in the analysis. Nevertheless, storing metadata in file names has some drawbacks, which will be addressed in future versions of monitoraSom.

### Template matching

Template matching is a technique used in passive acoustic monitoring (PAM) to detect specific sound patterns within large audio datasets by correlating recordings with acoustic templates containing the target species' sounds. When the structure of a template closely matches a segment of the soundscape, high correlation values (i.e., a correlation peak) indicate what could likely be an occurrence of the target species' acoustic signal. The validation app can then be used to assess results, refine the detection accuracy, and export data for further ecological analysis. While not as precise as human audio inspection, template matching is highly efficient for large-scale sound analysis, making it a valuable tool for bioacoustic research.

To run a template matching analysis, two distinct datasets are required: environmental sound recordings typically obtained in large quantities (the soundscapes) and at least one template, which is usually a valid high-fidelity recording of the target species used as a reference for detection.

Let's start by gathering the metadata of the soundscapes.

```
df_soundscapes <- fetch_soundscape_metadata(
  soundscapes_path = "./soundscapes", recursive = TRUE, ncores = 1
)
glimpse(df_soundscapes)
```

```
## Rows: 12
## Columns: 6
## $ soundscape_path        <chr> "./soundscapes/W54393S25597_20201104_170000.wav~
## $ soundscape_file        <chr> "W54393S25597_20201104_170000.wav", "W54431S256~
## $ soundscape_duration    <dbl> 59.99454, 59.99454, 59.99454, 59.99454, 59.9945~
## $ soundscape_sample_rate <int> 24000, 24000, 24000, 24000, 24000, 24000, 24000~
## $ soundscape_bitrate     <int> 16, 16, 16, 16, 16, 16, 16, 16, 16, 16, 16, 16
## $ soundscape_layout      <chr> "mono", "mono", "mono", "mono", "mono", "mono",~
```

Now, let's select the searching template files of *Basileuterus culicivorus*, as gathered in the first part of this exercise.

```
df_templates <- fetch_template_metadata(
    templates_path = "./templates",
    recursive = TRUE
)
glimpse(df_templates)
```

```
## Rows: 6
## Columns: 11
## $ template_path        <chr> "./templates/Bcu_1_004.236-004.742s_03.151-09.201~
## $ template_file        <chr> "Bcu_1_004.236-004.742s_03.151-09.201kHz_1024wl_7~
## $ template_name        <chr> "Bcu_1_004.236-004.742s_03.151-09.201kHz_1024wl_7~
## $ template_label       <chr> "Basileuterus culicivorus", "Basileuterus culiciv~
## $ template_start       <dbl> 0, 0, 0, 0, 0, 0
## $ template_end         <dbl> 0.5053750, 0.4963333, 0.5098750, 0.3642500, 0.400~
## $ template_sample_rate <int> 24000, 24000, 24000, 24000, 24000, 24000
## $ template_min_freq    <dbl> 3.151, 3.151, 3.057, 2.900, 2.975, 2.878
## $ template_max_freq    <dbl> 9.201, 9.232, 9.232, 9.764, 9.554, 9.636
## $ template_wl          <dbl> 1024, 1024, 1024, 1024, 1024, 1024
## $ template_ovlp        <dbl> 70, 70, 70, 70, 70, 70
```

By combining both data frames, we create a comprehensive map of correlations to be performed during the cross-correlation analysis.

```
df_grid <- fetch_match_grid(
    soundscape_data = df_soundscapes,
    template_data = df_templates
)
glimpse(df_grid)
```

```
## Rows: 72
## Columns: 17
## $ soundscape_path        <chr> "./soundscapes/W54393S25597_20201104_170000.wav~
## $ soundscape_file        <chr> "W54393S25597_20201104_170000.wav", "W54393S255~
```

```
## $ soundscape_duration    <dbl> 59.99454, 59.99454, 59.99454, 59.99454, 59.9945~
## $ soundscape_sample_rate <int> 24000, 24000, 24000, 24000, 24000, 24000, 24000~
## $ soundscape_bitrate     <int> 16, 16, 16, 16, 16, 16, 16, 16, 16, 16, 16, 16,~
## $ soundscape_layout      <chr> "mono", "mono", "mono", "mono", "mono", "mono",~
## $ template_path          <chr> "./templates/Bcu_1_004.236-004.742s_03.151-09.2~
## $ template_file          <chr> "Bcu_1_004.236-004.742s_03.151-09.201kHz_1024wl~
## $ template_name          <chr> "Bcu_1_004.236-004.742s_03.151-09.201kHz_1024wl~
## $ template_label         <chr> "Basileuterus culicivorus", "Basileuterus culic~
## $ template_start         <dbl> 0, 0, 0, 0, 0, 0, 0, 0, 0, 0, 0, 0, 0, 0, 0, 0,~
## $ template_end           <dbl> 0.5053750, 0.4963333, 0.5098750, 0.3642500, 0.4~
## $ template_sample_rate   <int> 24000, 24000, 24000, 24000, 24000, 24000, 24000~
## $ template_min_freq      <dbl> 3.151, 3.151, 3.057, 2.900, 2.975, 2.878, 3.151~
## $ template_max_freq      <dbl> 9.201, 9.232, 9.232, 9.764, 9.554, 9.636, 9.201~
## $ template_wl            <dbl> 1024, 1024, 1024, 1024, 1024, 1024, 1024, 1024,~
## $ template_ovlp          <dbl> 70, 70, 70, 70, 70, 70, 70, 70, 70, 70, 70, 70,~
```

At this stage, users can filter specific template-soundscape combinations to optimize processing or meet experimental design requirements. This is one of the most important checkpoints of the workflow, as it allows users to have granular control over the analysis. The resulting data frame has one row per template-soundscape pair along with their metadata, which includes the template spectrogram parameters that will be used to perform the cross-correlation analysis.

Let's make a quick check to ensure that the template and soundscape sample rates are compatible.

```
df_grid %>%
  select(soundscape_sample_rate, template_sample_rate) %>%
  distinct()
```

```
## # A tibble: 1 x 2
##   soundscape_sample_rate template_sample_rate
##                    <int>                <int>
## 1                  24000                24000
```

Let's also check that the overlap and window length are appropriate for the analysis. We expect template overlap to be 70% and window length to be 1024.

```
unique(df_grid$template_ovlp)
```

```
## [1] 70
```

```
unique(df_grid$template_wl)
```

```
## [1] 1024
```

Now that we have defined the grid and confirmed sample rate compatibility, we can proceed with the template matching analysis. This process performs cross-correlation by systematically sliding each template across the time axis of the soundscapes, as specified in our grid. Cross-correlation is a powerful signal processing technique that quantifies the similarity between two signals (in this case, the template and sections of the soundscape) at different time offsets. The resulting correlation values indicate how well the template matches the soundscape at each position, allowing us to identify potential occurrences of the target species' vocalizations.

Because this is the most time-consuming step of the analysis, users should evaluate whether their hardware can efficiently handle the computation before proceeding. In this example, we run a small test using 12 soundscapes and 6 templates, which gives us a grid of 72 template matching operations. It took approximately 17 seconds to complete using 6 cores of an M2 Apple processor, or 65 seconds using 6 cores of an Intel i7-10700 (4.800GHz) processor in Ubuntu 24.04. However, processing time can vary significantly depending on the computer's specifications. The results will be saved in a CSV file within the `"detections"` folder.

Let's make a time check to see how long it takes to run the template matching analysis with 6 cores. Note: check if you have 6 cores available in your computer before running the analysis. If you have fewer cores, you can change the number. Avoid using all cores, as it might slow down the analysis.

```
# check the number of cores available
# parallel::detectCores()

run_matching(
  df_grid = df_grid, output = "detections", ncores = 6,
  output_file = "detections/df_detecs.csv"
)
```

If results with parallel processing and single-core processing are not significantly different, parallelization is not working properly. There can be several reasons for this, as parallelization is handled differently across operating systems (MacOS, Windows, or Linux) and IDEs (RStudio, VSCode, etc.).

### Validation and template matching evaluation

The main objective of the validation process is to determine the optimal correlation threshold for efficient detection. The correlation threshold value directly influences the number of true positives and false positives in the results. While a higher correlation threshold tends to reduce false positives by ensuring that only strong matches are retained, it also reduces the number of true positives as the model becomes more selective. On the other hand, a lower correlation threshold increases recall by capturing more true positives, but at the cost of lower precision, as it leads to a higher number of false positives above the threshold. The ideal threshold strikes a balance between precision and recall, ensuring sufficient data for ecological analysis while maintaining the quality of the results.

We will first demonstrate the validation process using the validation app, within which users can validate the detections manually. Because validation takes place with data produced after the template matching process, this validation is referred to as *a posteriori* validation.

The file we will pass to the `input_path` argument should be the output of the `run_matching()` function, i.e., "detections/df_detecs.csv". However, it might be important to save the output file with a different name than the input file to preserve data and avoid rerunning the template matching process. With the steps below, we will make a copy of the detections file and use it as the input for the validation app.

```
original_file <- "detections/df_detecs.csv"
validation_file <- "validation_outputs/df_detecs_val_manual.csv"
# check if original file exists and copy doesn't exist yet
if (file.exists(original_file) && !file.exists(validation_file)) {
  file.copy(from = original_file, to = validation_file)
} else if (!file.exists(original_file)) {
  stop("The file 'detections/df_detecs.csv' does not exist.")
} else {
  message("Validation copy already exists, skipping copy.")
}
```

```
## Validation copy already exists, skipping copy.
```

Now we can launch the validation app. Note that we are using the validation copy as both the input file and the output file. This is a good practice to avoid overwriting the original data and to ensure we can validate the same data across multiple sessions.

```r
launch_validation_app(
  project_path = ".",
  validation_user = "Rosa G. L. M.",
  templates_path = "templates/",
  soundscapes_path = "soundscapes/",
  input_path = "validation_outputs/df_detecs_val_manual.csv",
  output_path = "validation_outputs/df_detecs_val_manual.csv",
  dyn_range_templ = c(-42, 0), dyn_range_detec = c(-90, -48),
  wl = 512, ovlp = 70, time_pads = 0.5, nav_autosave = TRUE, overwrite = TRUE,
  auto_next = TRUE, path_update = FALSE
)
```

Once we launch the Validation App, we can validate the detections as true or false positives and set a correlation threshold that balances precision and recall. Similar to the Segmentation App, the Validation App requires specific inputs to function properly. The project path is typically set to the working directory, and the user's name must be provided. Additionally, the paths to the template and soundscape files should be specified. Within the app, users must check the information and confirm the paths by clicking the 'confirm path' button. Users must then advance to the session setup tab, also in the sidebar menu, to confirm the validation setup. Within this menu, users can select which template to validate, the correlation score interval, or determine a number of the top scores to be validated. Additionally, the 'Order by' dropdown menu allows users to control the order in which the detections are presented for validation.

The app presents two spectrograms to facilitate the validation process. The spectrogram parameters can be adjusted through the spectrogram menu. The left spectrogram displays the detection from the soundscape file, while the right shows the template used in the correlation. Users can easily classify the detection as false (Q), positive (E), or unknown (W). If the Autosave and Autonavigate boxes are checked, the validation process becomes very efficient, automatically saving the classifications and navigating to the next detection. Additionally, users can listen to either the detection or the template to assist the validation process. As the validation process advances, the detection table is updated with the validation inputs, and the resulting data is used in a diagnostic analysis model employing a binomial regression. The validation app is typically set to assess correlation values with a precision of 95% (5% error). Along with the binomial regression plot, users can also see the density plots of correlation values for both false positives and true positives, and the precision vs recall plot. All these plots are updated in real-time, allowing users to evaluate the outcomes of the validation in real-time.

It should be noted that the only output of the validation app are the validation inputs routed to the CSV file passed to the `input_path` argument. The diagnostics data and plots are only displayed in the app, but can be computed with the `diagnostic_validations()` function. Let's import the validated detections and compute the diagnostics.

```r
df_detecs_val_manual <- read.csv(
  "validation_outputs/df_detecs_val_manual.csv", row.names = 1
)
ls_validations <- diagnostic_validations(
  df_validated = df_detecs_val_manual, pos_prob = 0.9
)
```

```
## Validation diagnostics completed successfully.
```

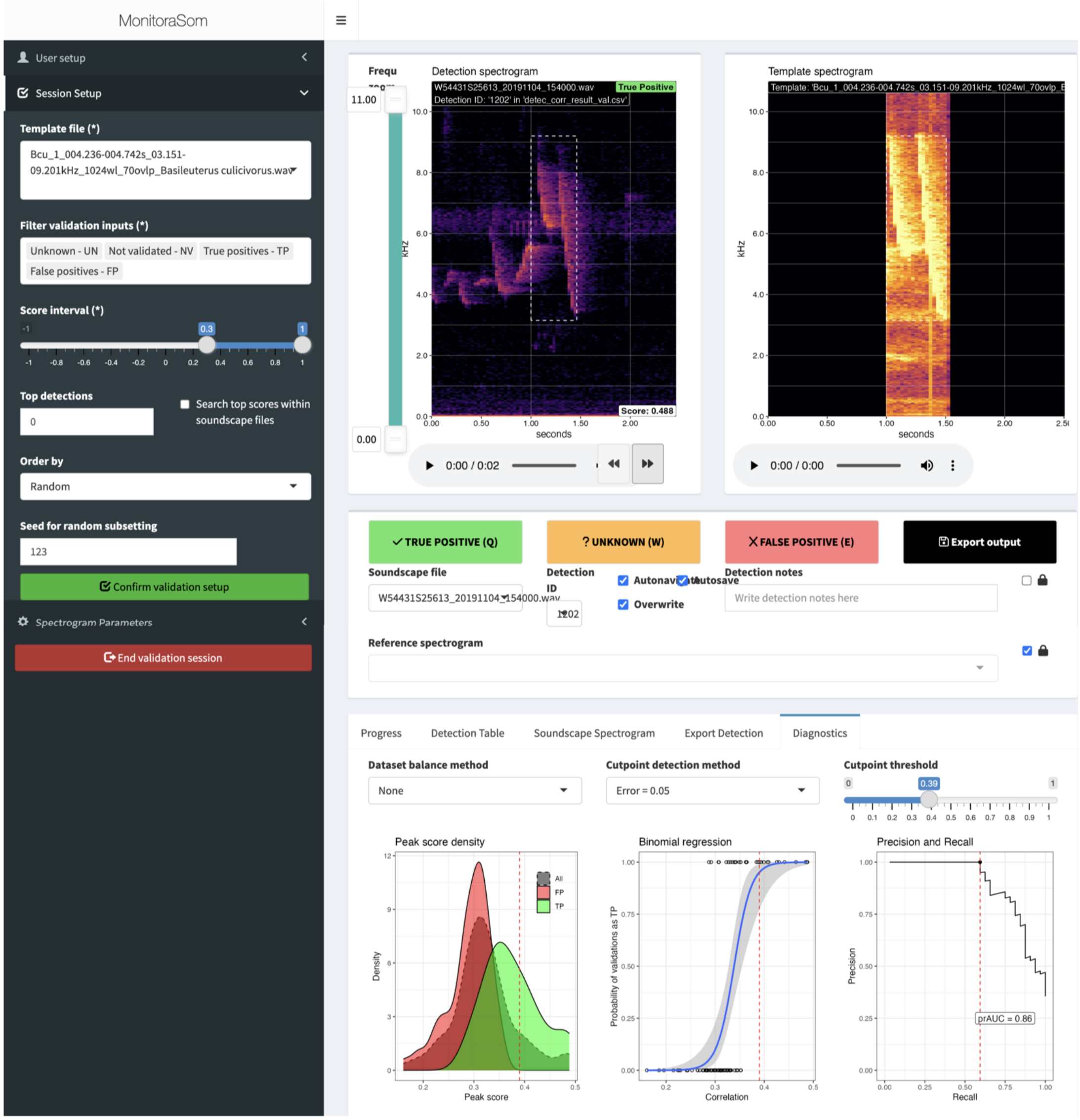


Figure 4: The validation app

Let's extract and plot the diagnostics for the first template. The `ls_validations` object is a list in which each element is a list of diagnostics for a given template. Now we can plot the diagnostics of the first template.

```
diags_template01 <- ls_validations[[1]]
diags_template01$mod_plot +
  diags_template01$precrec_plot +
  diags_template01$f1_plot +
  diags_template01$plot_dens
```

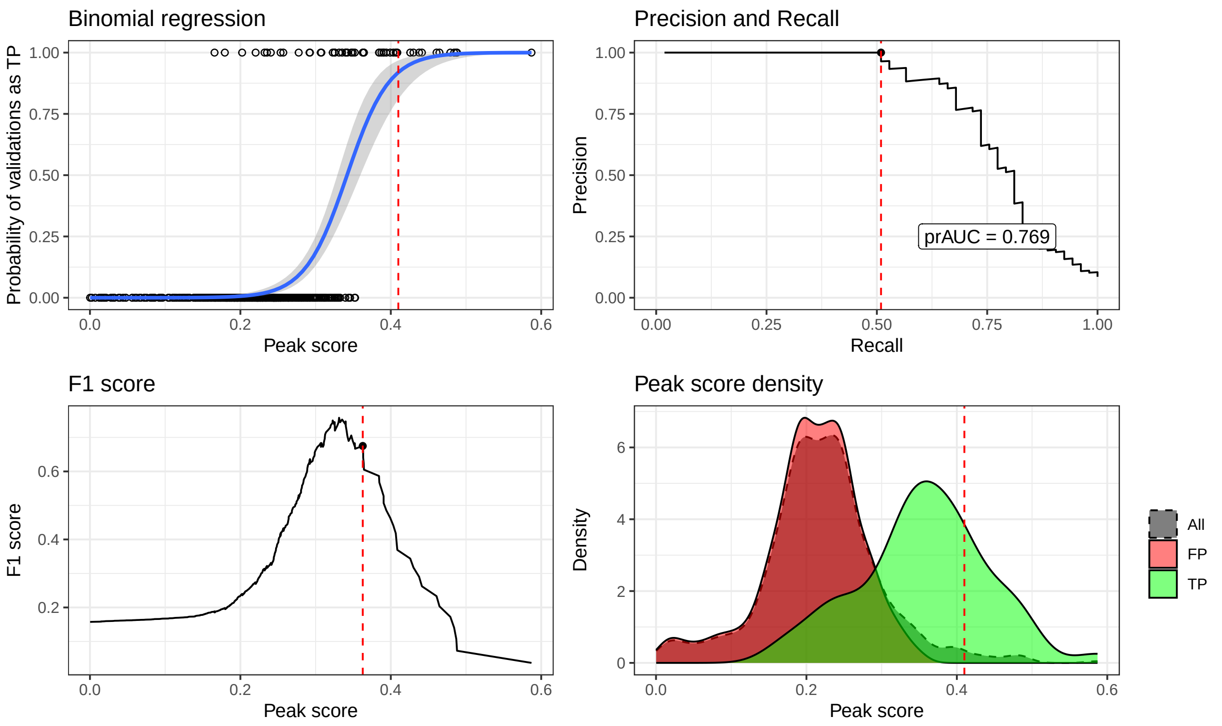


We can see that the top-left panel shows the plot of the binomial model used to estimate the score threshold with a minimum precision of 95% (5% error). It shows that we were able to estimate the score threshold with a minimum precision of 95% (5% error), and although no false positives were found, we missed many true positives (i.e., we have many false negatives). The top-right panel shows the precision-recall plot, which shows the precision and recall of the template matching process at different correlation thresholds. We can see in this case that we can reach a recall slightly above 50% with full precision, yet to increase the recall, we would need to accept lower precision. The bottom-left panel shows the F1 score plot, which shows the F1 score of the template matching process at different correlation thresholds. The F1 score is the harmonic mean of precision and recall, and it is a good indicator of the overall performance of the template matching process. Yet, because we will prioritize precision over recall, the most adequate threshold is the one that maximizes precision. The bottom-right panel shows the density plot of the correlation values for both false positives and true positives. The extensive tail of the false positives distribution shows that the template matching process is very good at detecting the target species, but it is not perfect.

Before extracting the final set of validations based on the correlation threshold, let's try another type of validation and compare the results. The alternative to the validation app is to use the `validate_by_overlap()` function, which allows users to validate the detections automatically based on the overlap between the detections and ROI tables of the soundscape recordings produced within the segmentation app. This validation method is referred to as *a priori* validation, as it is based on data previously available, namely the segmentation data from the soundscape recordings.

Before proceeding, we need to import the ROIs.

```r
df_rois <- fetch_rois(rois_path = "./roi_tables/")
glimpse(df_rois)
```

```
## Rows: 71
## Columns: 19
## $ soundscape_path      <chr> "./recordings/Bcu_1.wav", "./recordings/Bcu_1.wav~
## $ soundscape_file      <chr> "Bcu_1.wav", "Bcu_1.wav", "Bcu_1.wav", "Bcu_1.wav~
## $ roi_path             <chr> "/home/grosa/R_repos/monitoraSom/example/vignette~
## $ roi_file             <chr> "Bcu_1_roi_User_20250226101835.csv", "Bcu_1_roi_U~
## $ roi_user             <chr> "User", "User", "User", "User", "User", "User", "~
## $ roi_input_timestamp  <chr> "2025-02-26 10:19:57", "2025-02-26 10:20:12", "20~
## $ roi_label            <chr> "Basileuterus culicivorus", "Basileuterus culiciv~
## $ roi_start            <dbl> 4.236420, 15.691662, 27.506793, 3.794958, 15.2593~
## $ roi_end              <dbl> 4.741796, 16.188014, 28.016681, 4.189475, 15.6949~
## $ roi_min_freq         <dbl> 3.150896, 3.150896, 3.057099, 3.659274, 3.561569,~
## $ roi_max_freq         <dbl> 9.200792, 9.232058, 9.232058, 5.629660, 5.743649,~
## $ roi_type             <chr> "bird - song", "bird - song", "bird - song", "bir~
## $ roi_label_confidence <chr> "certain", "certain", "certain", "certain", "cert~
## $ roi_is_complete      <chr> "complete", "complete", "complete", "complete", "~
## $ roi_comment          <chr> "Substructure C", "Substructure C", "Substructure~
## $ roi_wl               <int> 1024, 1024, 1024, 1024, 1024, 1024, 1024, 1024, 1~
## $ roi_ovlp             <int> 70, 70, 70, 70, 70, 70, 70, 70, 70, 70, 70, 70, 7~
## $ roi_sample_rate      <int> 24000, 24000, 24000, 24000, 24000, 24000, 24000, ~
## $ roi_pitch_shift      <int> 1, 1, 1, 1, 1, 1, 1, 1, 1, 1, 1, 1, 1, 1, 1, 1, 1~
```

Now we can validate the detections by overlap with the ROIs. Detections that overlap with at least one ROI in the time window are validated as true positives, and all other detections are validated as false positives.

```r
df_detecs_val_tovlp <- validate_by_overlap(
  df_detecs = df_detecs, df_rois = df_rois, validation_user = "User Name",
  output_path = "validation_outputs/df_detecs_val_tovlp.csv"
)
```

```
## All detected species have ROIs for validation
```

```
## Validation results have been saved to validation_outputs/df_detecs_val_tovlp.csv
```

```r
glimpse(df_detecs_val_tovlp)
```

```
## Rows: 4,473
## Columns: 45
## $ soundscape_path        <chr> "soundscapes//Bcu_1.wav", "soundscapes//Bcu_1.wa~
## $ soundscape_file        <chr> "Bcu_1.wav", "Bcu_1.wav", "Bcu_1.wav", "Bcu_1.wa~
## $ template_path          <chr> "./templates//Bcu_1_004.236-004.742s_03.151-09.2~
## $ template_file          <chr> "Bcu_1_004.236-004.742s_03.151-09.201kHz_1024wl_~
## $ template_name          <chr> "Bcu_1_004.236-004.742s_03.151-09.201kHz_1024wl_~
## $ template_min_freq      <dbl> 3.151, 3.151, 3.151, 3.151, 3.151, 3.151, 3.151,~
## $ template_max_freq      <dbl> 9.201, 9.201, 9.201, 9.201, 9.201, 9.201, 9.201,~
## $ template_start         <int> 0, 0, 0, 0, 0, 0, 0, 0, 0, 0, 0, 0, 0, 0, 0, 0, ~
```

```
## $ template_end         <dbl> 0.505375, 0.505375, 0.505375, 0.505375, 0.505375~
## $ detection_start      <dbl> NA, NA, NA, NA, NA, NA, NA, NA, NA, NA, NA, NA, ~
## $ detection_end        <dbl> NA, NA, NA, NA, NA, NA, NA, NA, NA, NA, NA, NA, ~
## $ detection_wl         <int> NA, NA, NA, NA, NA, NA, NA, NA, NA, NA, NA, NA, ~
## $ detection_ovlp       <int> NA, NA, NA, NA, NA, NA, NA, NA, NA, NA, NA, NA, ~
## $ detection_sample_rate <int> NA, NA, NA, NA, NA, NA, NA, NA, NA, NA, NA, NA, ~
## $ detection_buffer     <int> NA, NA, NA, NA, NA, NA, NA, NA, NA, NA, NA, NA, ~
## $ detection_min_score  <lgl> NA, NA, NA, NA, NA, NA, NA, NA, NA, NA, NA, NA, ~
## $ detection_min_quant  <lgl> NA, NA, NA, NA, NA, NA, NA, NA, NA, NA, NA, NA, ~
## $ detection_top_n      <lgl> NA, NA, NA, NA, NA, NA, NA, NA, NA, NA, NA, NA, ~
## $ peak_index           <int> NA, NA, NA, NA, NA, NA, NA, NA, NA, NA, NA, NA, ~
## $ peak_score           <dbl> NA, NA, NA, NA, NA, NA, NA, NA, NA, NA, NA, NA, ~
## $ peak_quant           <dbl> NA, NA, NA, NA, NA, NA, NA, NA, NA, NA, NA, NA, ~
## $ species              <chr> "Basileuterus culicivorus", "Basileuterus culici~
## $ detection_id         <int> NA, NA, NA, NA, NA, NA, NA, NA, NA, NA, NA, NA, ~
## $ roi_path             <chr> "/home/grosa/R_repos/monitoraSom/example/vignett~
## $ roi_file             <chr> "Bcu_1_roi_User_20250226101835.csv", "Bcu_1_roi_~
## $ roi_user             <chr> "User", "User", "User", "User", "User", "User", ~
## $ roi_input_timestamp  <chr> "2025-02-26 10:19:57", "2025-02-26 10:20:12", "2~
## $ roi_label            <chr> "Basileuterus culicivorus", "Basileuterus culici~
## $ roi_start            <dbl> 4.236420, 15.691662, 27.506793, 3.794958, 15.259~
## $ roi_end              <dbl> 4.741796, 16.188014, 28.016681, 4.189475, 15.694~
## $ roi_min_freq         <dbl> 3.150896, 3.150896, 3.057099, 3.659274, 3.561569~
## $ roi_max_freq         <dbl> 9.200792, 9.232058, 9.232058, 5.629660, 5.743649~
## $ roi_type             <chr> "bird - song", "bird - song", "bird - song", "bi~
## $ roi_label_confidence <chr> "certain", "certain", "certain", "certain", "cer~
## $ roi_is_complete      <chr> "complete", "complete", "complete", "complete", ~
## $ roi_comment          <chr> "Substructure C", "Substructure C", "Substructur~
## $ roi_wl               <int> 1024, 1024, 1024, 1024, 1024, 1024, 1024, 1024, ~
## $ roi_ovlp             <int> 70, 70, 70, 70, 70, 70, 70, 70, 70, 70, 70, 70, ~
## $ roi_sample_rate      <int> 24000, 24000, 24000, 24000, 24000, 24000, 24000,~
## $ roi_pitch_shift      <int> 1, 1, 1, 1, 1, 1, 1, 1, 1, 1, 1, 1, 1, 1, 1, 1, ~
## $ roi_id               <int> 1, 2, 3, 4, 5, 6, 7, 8, 9, 10, 11, 12, 13, 14, 1~
## $ validation_user      <chr> "User Name", "User Name", "User Name", "User Nam~
## $ validation_time      <chr> "2025-06-23 15:46:35", "2025-06-23 15:46:35", "2~
## $ validation           <chr> "FN", "FN", "FN", "FN", "FN", "FN", "FN", "FN", ~
## $ validation_note      <chr> "no detections to intersect with", "no detection~
```

But if we count the number of detections by validation outcome, we see that there is a third outcome, which is the FN or false negatives. False negatives, in this case, are ROIs that were missed by the template matching process, i.e., ROIs that do not overlap with any detection.

```r
count(df_detecs_val_tovlp, validation)
```

```
## # A tibble: 3 x 2
##   validation     n
##   <chr>      <int>
## 1 FN           110
## 2 FP          3741
## 3 TP           622
```

If we perform the validation diagnostics again, we see that taking FN into account does interfere with the performance metrics of the template matching process. Lets compute the diagnostics again and plot the results.

```r
ls_validations_tovlp <- diagnostic_validations(
  df_validated = df_detecs_val_tovlp, pos_prob = 0.85
)
```

```
## Validation diagnostics completed successfully.
```

```r
diags_template01_tovlp <- ls_validations_tovlp[[1]]
diags_template01_tovlp$mod_plot +
  diags_template01_tovlp$precrec_plot +
  diags_template01_tovlp$f1_plot +
  diags_template01_tovlp$plot_dens
```

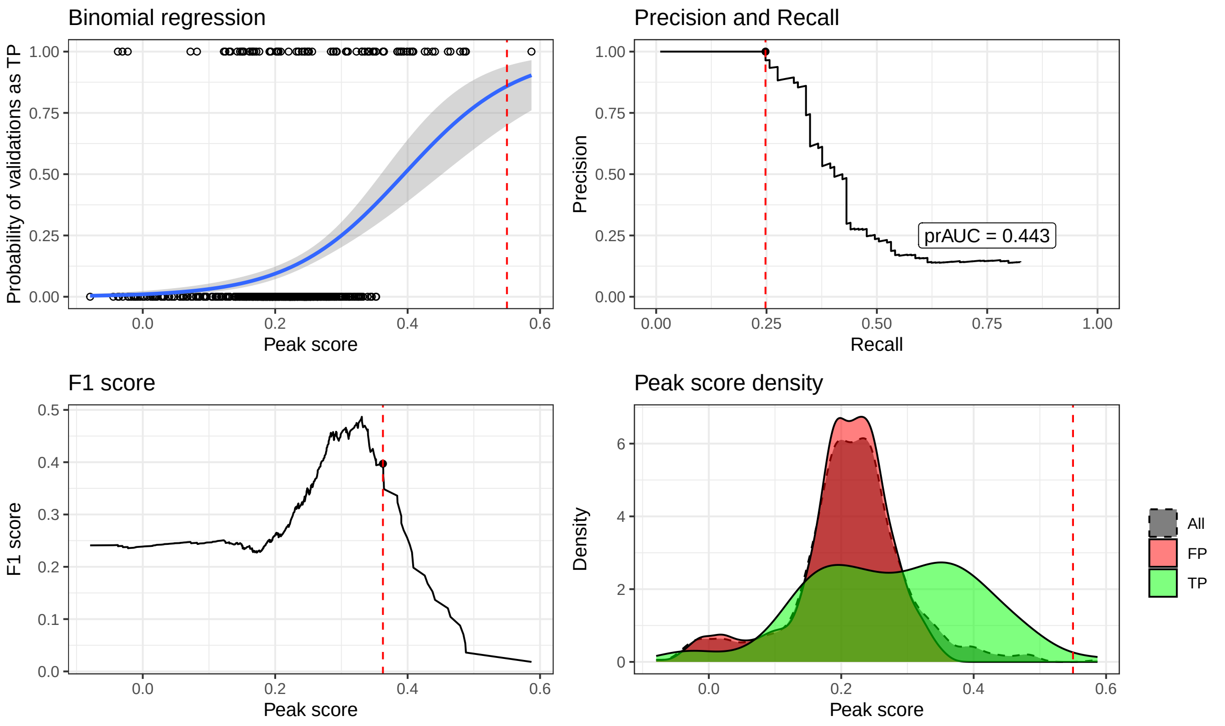


When comparing templates, users should look for templates with a clearly defined probability of classification as true in the binomial model, which sustain high precision and recall at the same time, with high F1 scores at the selected precision threshold and a clear separation between the TP and FP distributions. A closer look into the diagnostics of the second round of validation (*a priori*) shows a few important patterns. First, we had to lower the precision threshold to 0.85 to stay within the testable range of the binomial model. We also see that precision starts to drop steeply with recalls above 25%, which is much sooner than the 50% of the *a posteriori* approach. The compromises between precision and recall are similar to the *a posteriori* validation, but the F1 score range is lower, which shows a worse overall detection performance. The density plot of the TP and FP distributions shows a less clear separation between the two distributions.

The soundscapes, templates, and detections are the same as in the *a posteriori* validation, but there as fundamenetal differences between the two validation methods.Because the *a posteriori* validation does not account for false negatives, we see that although the template is useful, maybe we should keep looking for a better template. Commenting further on the *a posteriori* validation, detection tables that will be validated with this approach should not be heavily filtered, as it may result in artificial inflation of FN counts. This is why we stress that ROI tables used for *a posteriori* validation must be complete (i.e., include all signals from the target species in the respective soundscape recording). As a rule of thumb, we recommend using the validation app for exploratory analyses and the validation by overlap for large-scale use cases.

Sometimes it is not easy to find a template that satisfies all criteria mentioned above, which is why we recommend always evaluating a set of different templates under varying settings. But, for now, we are satisfied with the results, then we can export the final set of validations based on the correlation threshold. Here we demonstrate a few steps to ensure we export the detections based on the correct template and threshold.

```r
# Let's get the results of the first template to ensure consistency
res <- ls_validations_tovlp[[1]]

# first we retrieve the threshold determined by the diagnostics
selected_thres <- res$score_cut

# then we extract the template name from the diagnostics to ensure we are
# filtering the detections based on the correct template
first_template_name <- unique(res$diagnostics$template_name)

# Then we simply filter the detections based on the template name and the
# threshold value for that template
df_detecs_final <- df_detecs_val_manual %>%
  filter(template_name == first_template_name, peak_score >= selected_thres) %>%
  glimpse()
```

```
## Rows: 1
## Columns: 26
## $ soundscape_path       <chr> "./soundscapes//W54448S25622_20191104_171000.wav"
## $ soundscape_file       <chr> "W54448S25622_20191104_171000.wav"
## $ template_path         <chr> "./templates//Bcu_1_004.236-004.742s_03.151-09.2~
## $ template_file         <chr> "Bcu_1_004.236-004.742s_03.151-09.201kHz_1024wl_~
## $ template_name         <chr> "Bcu_1_004.236-004.742s_03.151-09.201kHz_1024wl_~
## $ template_min_freq     <dbl> 3.151
## $ template_max_freq     <dbl> 9.201
## $ template_start        <int> 0
## $ template_end          <dbl> 0.505375
## $ detection_start       <dbl> 57.11204
## $ detection_end         <dbl> 57.57324
## $ detection_wl          <int> 1024
## $ detection_ovlp        <int> 70
## $ detection_sample_rate <int> 24000
## $ detection_buffer      <int> 37
## $ detection_min_score   <lgl> NA
## $ detection_min_quant   <lgl> NA
## $ detection_top_n       <lgl> NA
## $ peak_index            <int> 4477
## $ peak_score            <dbl> 0.5870497
## $ peak_quant            <dbl> 1
## $ detection_id          <int> 3701
## $ validation_user       <lgl> NA
## $ validation_time       <chr> "2025-03-06 10:49:02"
## $ validation            <chr> "TP"
## $ validation_note       <lgl> NA
```

Although in this example we retrieved a final set of one detection above the threshold for this template, this was retrieved from a single template in a significantly smaller set of soundscapes. Use cases in larger set of soundscape recordings would return a much larger quantity of useful detections. Also, results from multiple templates can be combined to achieve both increased performance and complementarity, leading to a more robust set of detections.

With this we desmontrated the basic workflow of the template matching process and the next steps are to prepare the detections for ecological analysis.

#### Summary

*A typical template matching analysis is made by the following steps:*

1. *Data Collection.* Gather a large dataset of audio recordings, typically from soundscapes that include the target species' calls or sounds.
2. *Template Creation.* Define or select acoustic templates that represent the target species' sounds. These templates are usually manually curated from high-fidelity recordings and represent the characteristic frequency patterns of the species' calls.
3. *Preprocessing of Audio.* Process the audio data to prepare it for analysis. This can include editing the audio recordings to enhance the signal-to-noise ratio of the templates, segmenting audio into manageable parts, or changing the sampling rate of the template so that it matches the sampling rate of the soundscapes.
4. *Fetching templates.* Select the templates and include them in an R object.
5. *Fetching Soundscapes.* Select the soundscapes folder and create a list in an R object.
6. *Fetching Soundscapes.* Create a correlation grid for the analysis.
7. *Template Matching.* Use the 'run_matching()' function to compute the correlations between the template and the soundscape at different time points.
8. *Thresholding and Identification.* Validate the detections to set a correlation threshold that balances precision and recall.
9. *Scale up.* Use the correlation threshold to filter out false positives from a larger set of soundscape recordings.
10. *Post validation.* Select a sample of the large-scale data to execute a final validation.
11. *Hypothesis testing.* Once the analysis is validated, export the results for further ecological analysis, such as determining species occupancy, activity patterns, or habitat use.
